# Compartment-Specific Measurement of Small Molecule Accumulation into Diderm Bacteria

**DOI:** 10.1101/2022.05.05.489932

**Authors:** George M. Ongwae, Irene Lepori, Mahendra D. Chordia, Brianna E. Dalesandro, Alexis J. Apostolos, M. Sloan Siegrist, Marcos M. Pires

## Abstract

Some of the most dangerous bacterial pathogens (Gram-negative and mycobacterial) deploy a formidable secondary membrane barrier to reduce the influx of exogenous molecules. For Gram-negative bacteria, this second exterior membrane is known as the outer membrane, while for the Gram-indeterminate *Mycobacteria,* it is known as the ‘myco’ membrane. Although different in composition, both the outer membrane and mycomembrane are key structures that restrict the passive permeation of small molecules into bacterial cells. While it is well-appreciated that such structures are principal determinants of small molecule permeation, it has proven to be challenging to assess this feature in a robust and quantitative way or in complex, infection-relevant settings. Herein, we describe the development of the <u>B</u>acterial <u>C</u>hloro-<u>A</u>lkane <u>P</u>enetration <u>A</u>ssay (BaCAPA), which employs the use of a genetically encoded protein called HaloTag, to measure the uptake and accumulation of molecules into model Gram-negative and mycobacterial species, *Escherichia coli* and *Mycobacterium smegmatis,* respectively, and into the human pathogen *M. tuberculosis.* Directing the localization of the HaloTag protein to either the cytoplasm or periplasm of bacteria enabled a compartmental analysis of permeation across individual cell membranes. Significantly, we also showed that BaCAPA can be used to analyze the permeation of molecules into host cell-internalized *E. coli* and *M. tuberculosis,* a critical capability for analyzing intracellular pathogens. Together, our results show that BaCAPA affords facile, compartment-specific measurement of permeability across four barriers: the host plasma and phagosomal membranes and the diderm bacterial cell envelope.

## Introduction

The rise in incidence of drug-resistant bacterial infections poses a tremendous challenge for healthcare systems throughout the world.^*1–3*^ While recent efforts have led to the discovery of new therapeutic leads with promising clinical potential,^*4–10*^ there is a continued need to strengthen the antibiotic pipeline and to find antibiotics with narrow spectrum activities to reduce off target impact on gut commensal bacteria. Most antibiotics must enter the bacterial cell to impart their biological effects, which includes permeating through a lipid bilayer. For Gram-negative and mycobacterial species, there is the additional challenge due to a second, asymmetrical bilayer known as the outer membrane (OM)^*11,12*^ or the outer ‘myco’ membrane, ^*13*^ respectively. For some agents (e.g., β-lactam antibiotics), permeation through Gram-negative OM-anchored porins can potentially provide an access pathway to the periplasm.^*14*^ The human pathogen *Mycobacterium tuberculosis* lacks Gram-negative-like porins, however their PE/PPE proteins may serve analogous roles in nutrient and antibiotic uptake.^*15–19*^ Many of the most potent antibiotics have cytosolic targets, and, therefore to be effective they must permeate through not only external membranes, but also the inner plasma membrane. Additionally, a reduction in the amount of antibiotics accumulating intracellularly can be further modulated by the active removal of drugs that reach the periplasm by efflux pumps that recognize broad structural motifs.^*13, 20–23*^

Despite the fact that drug permeation is well recognized as a major bottleneck for antibiotic efficacy and directly tied to drug discovery efforts,^*24, 25*^ there are few methods developed to measure the accumulation of small molecules in bacteria.^*26*^ As such, gaps remain in our fundamental understanding of the molecular determinants of permeability. Direct methods of measuring drug uptake and retention can be extremely valuable in providing insight into drug accumulation profiles. The Hergenrother group^*27*^ adopted a protocol^*28*^ that uses liquid chromatography with tandem mass spectrometry (LC-MS/MS) to expand the analysis of drug permeation across 180 diverse molecules in *Escherichia coli.* A principal finding from these investigations was that primary amines as a functional group are privileged moieties in promoting accumulation in Gram-negative bacteria. More recent studies leveraged biorthogonal reactions to probe the accumulation to distinct compartments within *E. coli*^*29,30*^ or microspectrofluorimetry.

To date, most bacterial permeability studies have focused on *E. coli* as a model organism. However, it is unlikely that predictive rules for small molecule accumulation for *E. coli* suffice for mycobacterial species. For example, the overall permeability of the pathogenic non-tuberculous mycobacterium (NTM) *M. chelonae* to small, hydrophilic compounds is estimated to be ~10-1000-fold less than Gram-negatives.^*34,35*^ A study of 10 sulfonyladenosines also suggested that permeation rules in the model mycobacterium *M. smegmatis* may be different than those for *E. coli*.^*28*^ Further, the *M. tuberculosis* envelope is remodeled and efflux pump gene expression changes upon host-mimicking stress and during infection^*13, 36–39*^, changes that can correlate with dramatic differences in drug susceptibility. There is a significant gap in our understanding of mycobacterial permeability, particularly in infection-relevant context.

We sought to complement existing strategies for measuring bacterial permeability by retaining the targeted compartmentalization of detection while simplifying the mode of accumulation readout. More specifically, we adapted for diderm bacteria the chloroalkane penetration assay (CAPA) developed by the Kritzer lab^*40*^, which is now widely used to measure the accumulation of molecules into mammalian cells. This method relies on the genetically encoded HaloTag^*41*^, a mutant form of a bacterial haloalkane dehalogenase that facilitates covalent bond formation with chloroalkane-linked ligands (**Figure 1**). Previous efforts utilizing the HaloTag protein within bacteria focused on the visualization of protein fusions^*42, 43*^ and the role of a cationic peptide on membrane permeabilization.^*44*^ Building on these precedents, we developed the <u>B</u>acterial <u>C</u>hloroalkane <u>P</u>enetration <u>A</u>ssay (BaCAPA) to measure the apparent accumulation of molecules into bacterial cells. If the molecules reach the HaloTag proteins inside the bacteria, they will be covalently bound to the protein. This pulse step is followed by a chase step with a fluorophore-linked chloroalkane. In the absence of drug accumulation, HaloTag active sites remain available to react with the fluorophore. Conversely, when a large fraction of molecules of interest permeate the cell membranes and reach the HaloTag proteins, there will be reduced sites available to react with the fluorophore. Therefore, a decrease in cellular fluorescence signal should reflect the apparent drug accumulation into bacterial cells. BaCAPA readily and robustly measured small molecule accumulation in *E. coli* and *M. tuberculosis* in broth or within host cells. Our ultimate goal is to establish structural determinants that favor permeation of small molecules across the cellular envelope of diderm bacteria *in vivo* to discover privileged antibiotic scaffolds.

**Figure 1.**
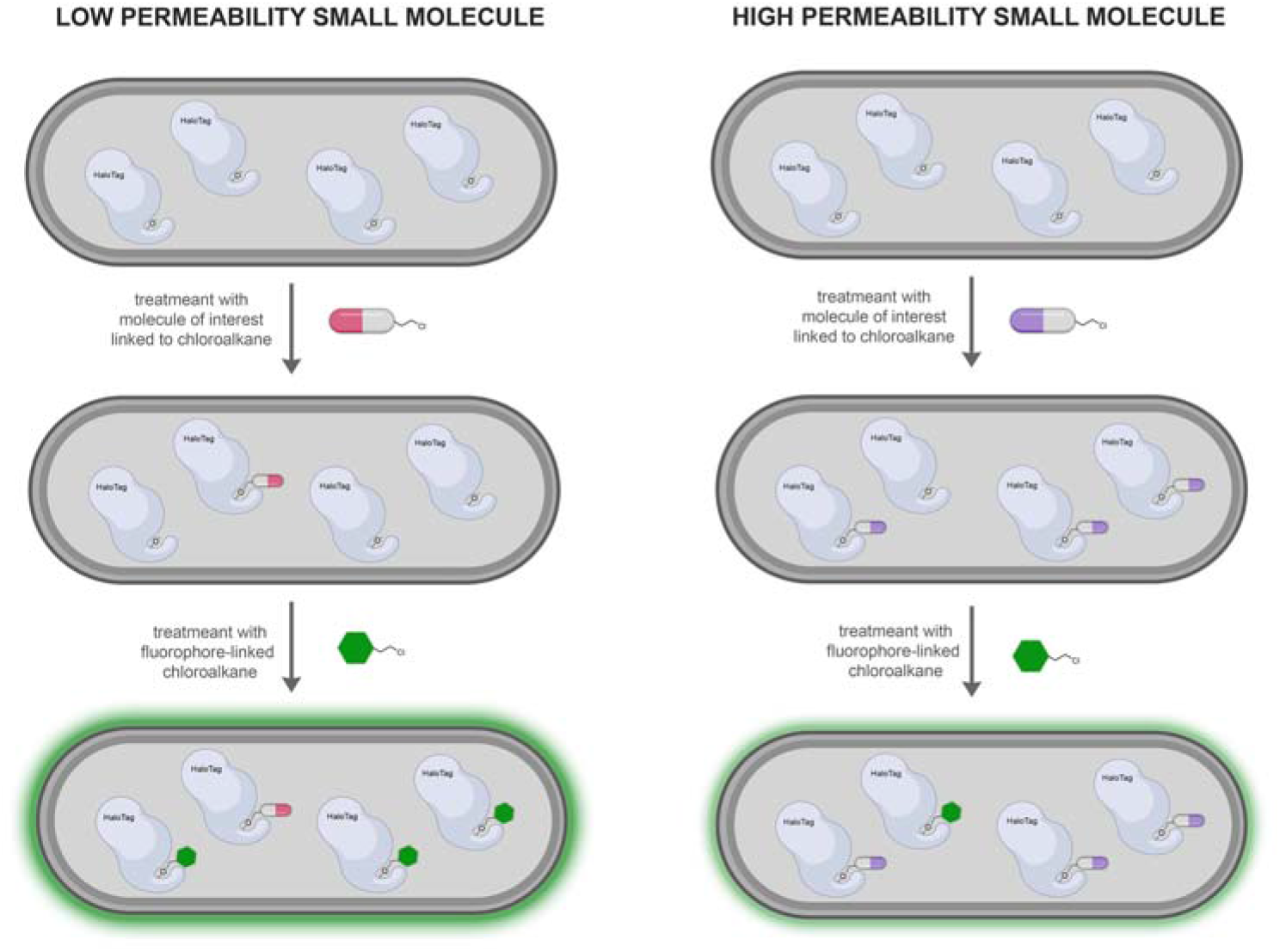
Schematic representation of BaCAPA. Bacterial cells expressing HaloTag will covalently react with molecules of interest, thus reducing the number of sites available to react with the fluorophore.

## Results

At first, we aimed to evaluate the fluorescence levels of *E. coli* that express HaloTag proteins in the cytosol by encoding the protein on an inducible plasmid. Bacterial cells carrying the HaloTag-expressing plasmid were grown to mid-log, induced with isopropyl-β-D-1-thiogalactopyranoside (IPTG), and incubated with a fluorophore-modified chloroalkane. Cytosolic HaloTag should promote the formation of a covalent bond with the fluorophore-linked chloroalkane tail (**Figure 2A**). Total labeling levels of bacterial cells can then be quantified using flow cytometry. To establish the first step in the BaCAPA, bacterial cells were treated with Rhodamine 110 modified with chloroalkane (**R110cl**). As expected, a large increase in cellular fluorescence was observed in a dose-dependent manner with increasing levels of IPTG (**Figure 2B**). These results suggest that HaloTag can report on the accumulation of **R110cl** into *E. coli.* Appreciating that the fluorophore itself could be subject to the same barrier elements that are inherent to *E. coli,* we also evaluated the labeling of bacterial cells with a smaller fluorophore, namely coumarin (**COMcl**). We projected that the physiochemical properties of these dyes could alter permeation into bacterial cells due to differences in size and charge between the two dyes. While it was clear that coumarin could also covalently modify HaloTag-expressing bacterial cells, the relative fluorescence increase was lower in cells treated with **COMcl** (**Figure 2C**) than with **R110cl**. Therefore, **R110cl** was chosen as the reporter dye for all subsequent studies in *E. coli*.

**Figure 2.**
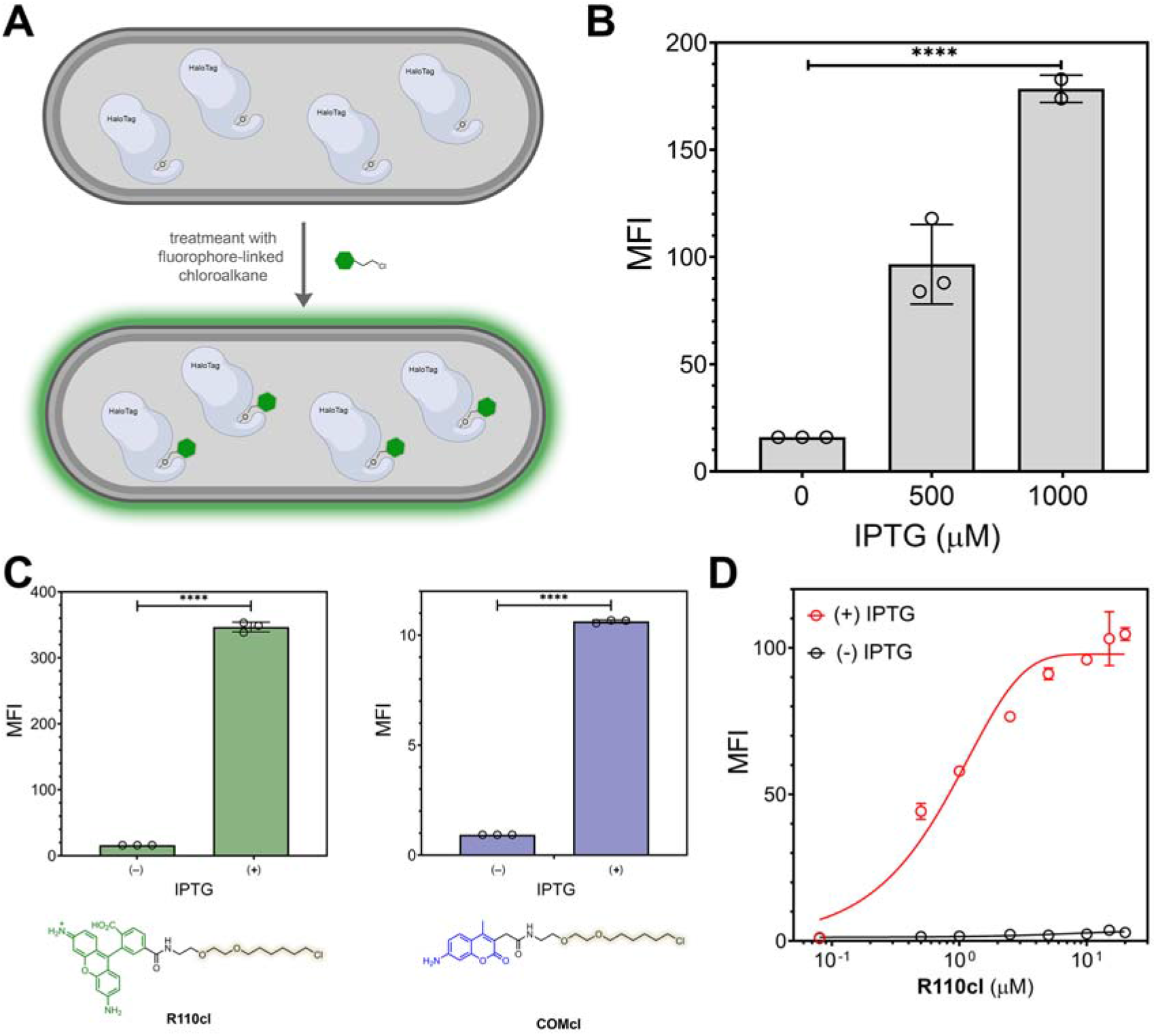
(A) Schematic representation of reaction of a fluorophore-linked to chloroalkane with HaloTag inside a bacterial cell. (B) Flow cytometry analysis of *E. coli* treated with various concentrations of IPTG, followed by an incubation period with 5 μM of **R110cl**. (C) Flow cytometry analysis of *E. coli* treated with either 5 μM of **R110cl** or **COMcl**. (D) Flow cytometry analysis of *E. coli* treated with various concentrations of **R110cl** in the presence and absence of IPTG induction. Data are represented as mean +/− SD (n = 3). *P*-values were determined by a two-tailed *t*-test (* denotes a *p-* value < 0.05, **□<□Q.01, ***<0.001, ns□=□not significant).

To empirically establish the dye concentration that optimizes the signal output for the assay, bacterial cells were treated with a range of **R110cl** concentrations (**Figure 2D**). The results showed that 0.5 μM is sufficient to yield fluorescence levels that were nearly 50% of the maximum and by 5 μM the fluorescence signals were reaching saturation levels. For these reasons, all subsequent assays were carried out using 5 μM of **R110cl**. We posited that it should be possible to analyze HaloTag labeling with **R110cl** from the whole cellular proteome because of the selectivity of this enzymatic reaction. To this end, bacterial cells were treated similar to the previous experiments, subjected to separation on an SDS-PAGE, and the gel was imaged using a fluorescent filter. As expected, there is a clear band that corresponds to the molecular weight of HaloTag (**Figure S1**), which strongly indicate that the signals observed from the flow cytometry analysis are representative of HaloTag covalently bound with **R110cl** inside bacterial cells. Cells were also imaged using confocal microscopy and the labelling pattern was consistent with cytoplasmic localization of HaloTag, as expected (**Figure S2**). Together, these results serve to establish the working parameters for BaCAPA.

It is well appreciated that the OM is the major barrier to the permeation of small molecules in Gram-negatives.^*45*^ Considerable efforts have been devoted to the discovery of molecules that disrupt the OM barrier as antibiotic adjuvants with the goal of potentiating them and/or circumventing resistance.^*46–49*^ Such agents hold the promise of synergistically enhancing the permeation of potential antibiotics in a broad and, potentially, impactful way. This class of molecules is best represented by the Gramnegative specific antibiotic polymyxin B. More specifically, a fragment called polymyxin B nonapeptide (PMBN), which lacks the fatty acid tail, has permeabilizing properties at concentrations that are non-toxic to bacterial cells.^*50*^ The potential clinical utility of OM permeabilizers is highlighted by the findings that PMBN protects mice against Gramnegative bacteria when dosed with erythromycin.^*51*^ We reasoned that co-treatment of cells with PMBN and **R110cl** should introduce a greater level of **R110cl** to bacterial cells, thus resulting in higher fluorescence levels (**Figure 3A**). Indeed, a concentrationdependent increase in cellular fluorescence was observed and the fluorescence levels of cells treated with 10 μM of PMBN were more than double that of cells not treated with PMBN. Moreover, no loss of bacterial viability was observed at the concentrations of PMBN tested (**Figure S3**).

**Figure 3.**
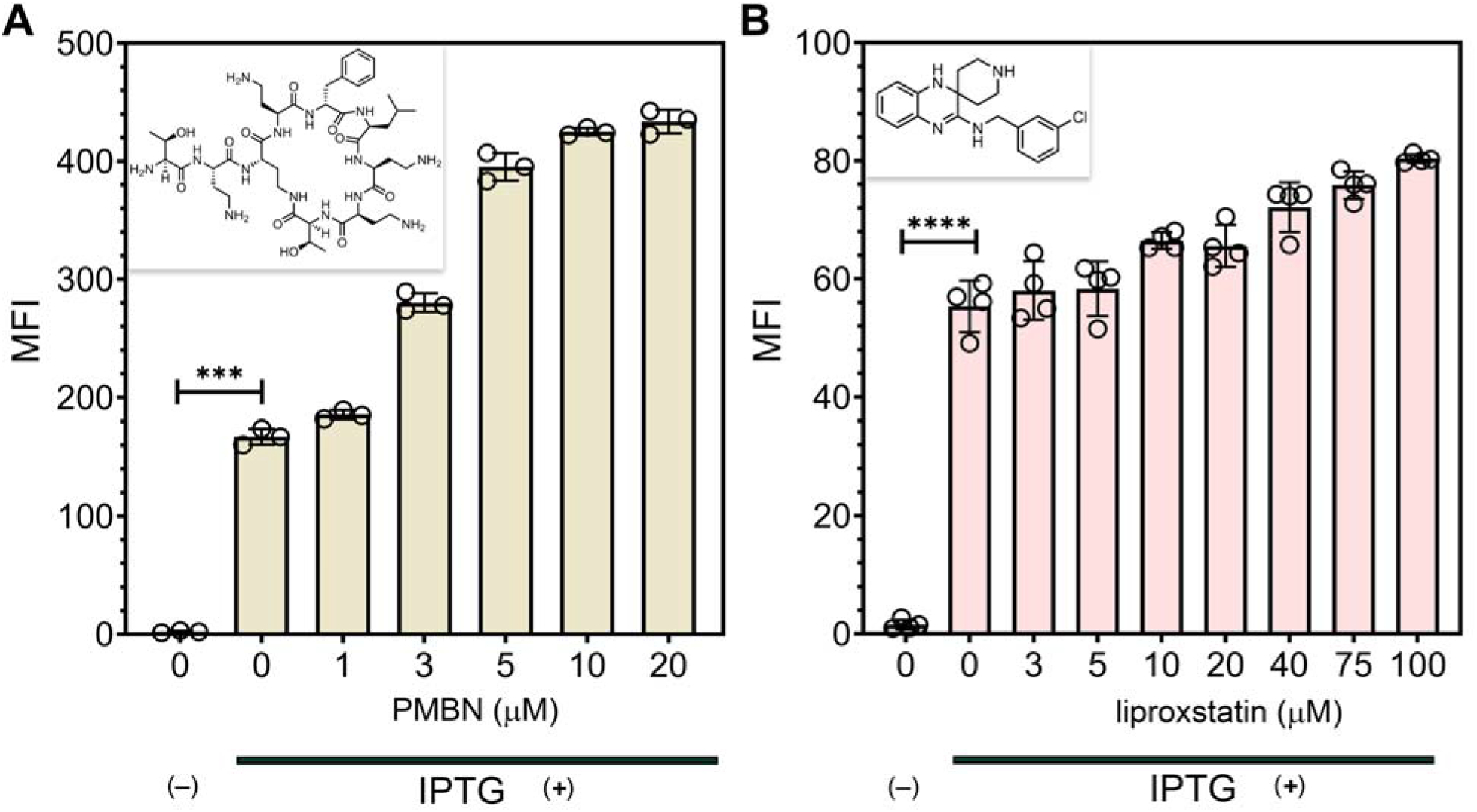
Flow cytometry analysis of *E. coli* treated with agents that are known to destabilize the outer membrane. After IPTG induction, *E. coli* cells were incubated with increasing concentrations of PMBN (A) or liproxstatin (B) for 30 min, washed, then treated with 5 μM of **R110cl**. Data are represented as mean +/− SD (n = 3). *P*-values were determined by a two-tailed *t*-test (* denotes a *p*-value < 0.05, **□<□0.01, ***<0.001, ns□=□not significant).

A recent screening campaign by the Brown laboratory against a library of 140,000 diverse compounds revealed liproxstatin as a promising new disruptor of *E. coli* OM.^*47*^ Similar to PMBN, an increase in cellular fluorescence was observed when liproxstatin was co-incubated with **R110cl** (**Figure 3B**). These results demonstrate the potential of BaCAPA to be leveraged, more generally, to report on molecules that disrupt the OM of Gram-negative bacteria. In an effort to test the assay compatibility with high-throughput screening platforms, we evaluated whether the cellular fluorescence levels could be measured *via* a plate reader rather than flow cytometer. Our results showed that the assay could readily report on HaloTag-mediated fluorescence (**Figure S4**). Our results showed that the signal strength in the presence of IPTG was well above that in the absence of IPTG. We are currently exploring the full compatibility of the assay to be used to screen for molecules that increase cellular permeability in Gram-negative bacteria.

Next, we set out to test whether BaCAPA could be implemented in a smooth *E. coli* strain (ATCC 25922) that contains the complete set of lipopolysaccharides on the surface. In contrast to K-12 rough bacteria, smooth *E. coli* strains can potentially have altered permeability. We recently demonstrated that the surface of rough *E. coli* had higher accessibility to molecules than smooth *E. coli,* presumably due to the steric blockade provided by *O*-antigens.^*52*^ Similarly, it is possible that the *O*-antigens may contribute to the overall permeation profile of small molecules considering that LPS chains have been proposed to occlude porin channels via/by steric shielding.^*53*^ Therefore, we set out to establish the feasibility of analyzing BaCAPA in smooth WT *E. coli* in a similar manner to that of the K-12 *E.* coli. When induced and non-induced WT *E.coli* cells were treated with **R110cl**, we observed a HaloTag dependent increase in fluorescence; therefore, these results confirmed that the permeability assay can be readily implemented in a WT strain (**Figure S5**). Additionally, we posed that it may be possible to take advantage of directed localization tags to specifically probe the accumulation of molecules within individual compartments in *E. coli.* To test this, an *N*-terminal fusion of a DsbA signal peptide was added to HaloTag to transport it to the periplasmic space.^*54*^ While we observed minimal fluorescence signals in *E. coli* BL21(DE3), high fluorescence levels were observed in Lemo21 (DE3) (**Figure S6**). Expression in Lemo21 cells likely afforded a more controlled expression level that may have favorably modulated the folding of HaloTag. Additionally, confocal microscopy established that the DsbA-HaloTag labeling pattern is more consistent with the expected localization within the periplasmic space compared to that of the cytoplasmic HaloTag (**Figure S7**). Fluorescence levels were found to be stronger on the periphery of the cells expressing DsbA-HaloTag.

We then set out to test the ability of BaCAPA to report on apparent accumulation of small molecules past the cellular envelope. In this format, (non-fluorescent) small molecules modified with the chloroalkane are incubated with the bacterial cells expressing HaloTag first (pulse step), which is then followed by a treatment step with the dye **R110cl** (chase step). The initial test was performed with a simple amino-ligand (**1**). A concentration scan of the small molecule revealed that nearly background fluorescence levels were observed at 12 μM with an EC_50_ ~ 5 μM (**Figure 4A**). These results suggest that BaCAPA can robustly and efficiently report on the apparent accumulation of small molecules into *E. coli.* We next wanted to evaluate how subtle changes to the chemical structure of the chase molecule could potentially impact its permeability into the bacterial cells. We decided to test how BaCAPA can be leveraged to evaluate the impact of methylation of a primary amine on relative accumulation in *E. coli* (**Figure 4B**). The Hergenrother lab previously demonstrated that the inclusion of a primary amine resulted in higher levels of accumulation in *E. coli* relative to any other methylation state across a series of molecules (although there were some deviations observed).^*27*^ Interestingly, our data revealed that within our series of molecules, there was minimal difference in relative accumulation across the primary, secondary, or tertiary amino group (compounds **2-4**) (**Figure 4B**). There was, however, an observed and statistically significant lowered accumulation level with the quaternary ammonium derivative (**5**). We pose that several factors could potentially explain our observations. First, the mode of permeation is likely to be an important molecular determinant towards the accumulation into the bacterial cells and its subsequent potential efflux. Therefore, the **2-4** series could be entering through a different route than prior molecules investigated by the Hergenrother group. Alternatively, we reason that the series chosen here has the potential for an internal hydrogen bond,^*55*^ which may alter the desolvation energetics, ultimately impacting permeation profiles across low dielectric environments.

**Figure 4.**
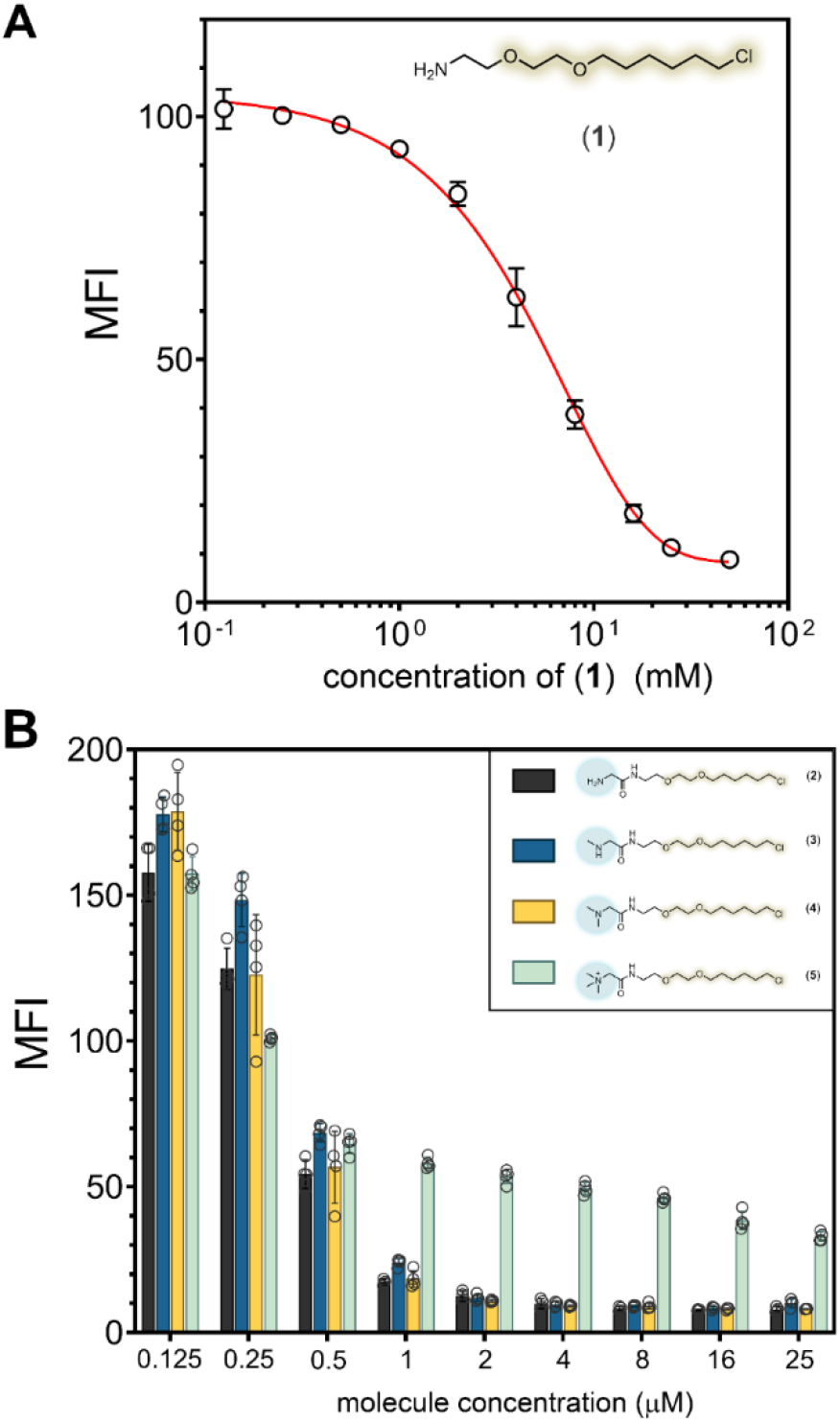
Flow cytometry analysis of *E. coli* treated with increasing concentrations of the canonical chloroalkane linker (**A**) or varying alkylation levels in molecules **2-5** (**B**), followed by a treatment with 5 μM of **R110cl** in the presence and absence of IPTG induction. Data are represented as mean +/− SD (n = 3).

To expand the application of BaCAPA for studying the permeability in *E. coli,* a group of molecules were synthesized based on the antibiotic ciprofloxacin decorated with a chloroalkane tag. After the discovery of quinolones as potent inhibitors of DNA replication in bacteria, this molecular scaffold was intensely explored in search of more potent analogs, which yielded clinically important agents like norfloxacin, sparfloxacin, levofloxacin, moxifloxacin, gatifloxacin.^*56*^ A large fraction of ciprofloxacin analogs involve derivatives to the piperazine moiety, which has shown to be tolerant of functional group manipulations. Therefore, we made two analogs that were built from modifications of the secondary amine for installation of the chloroalkane linker (**Figure 5**). In one analog, the secondary amine is converted to an amide (**6**) and another where the modification was made with the goal of retaining an ionizable amino group (**7**). A third molecule was synthesized in which the chloroalkane was, instead, coupled onto the carboxylic acid on the parent molecule (**8**). The free carboxylic acid is known to be crucial for the antimicrobial activity of ciprofloxacin, and, therefore, we did not expect that this molecule would have biological activity.^*57*^ Nonetheless, it provided an opportunity to assess how the permeation could be modulated by making a derivative that should have a net positive charge. Our results showed that retention of an ionizable amine group in (**7**) resulted in improved apparent permeation (EC_50_ = 6.5 μM) relative to the similarly structured analog with an amide (**6**), which had an EC_50_ = 42.2 μM (**Figure 5**). Interestingly, we found that (**6**) and (**7**) had the same MIC value of 0.5 μg/mL. These results highlight a complication that can arise from using MIC values as a proxy for relative permeation of molecules into Gram-negative bacteria. Installment of the chloroalkane ligand in (**8**) led to a significant reduction in EC_50_ (0.5 μM); nonetheless, the MIC value increased 8-fold to 4 μg/mL.

**Figure 5.**
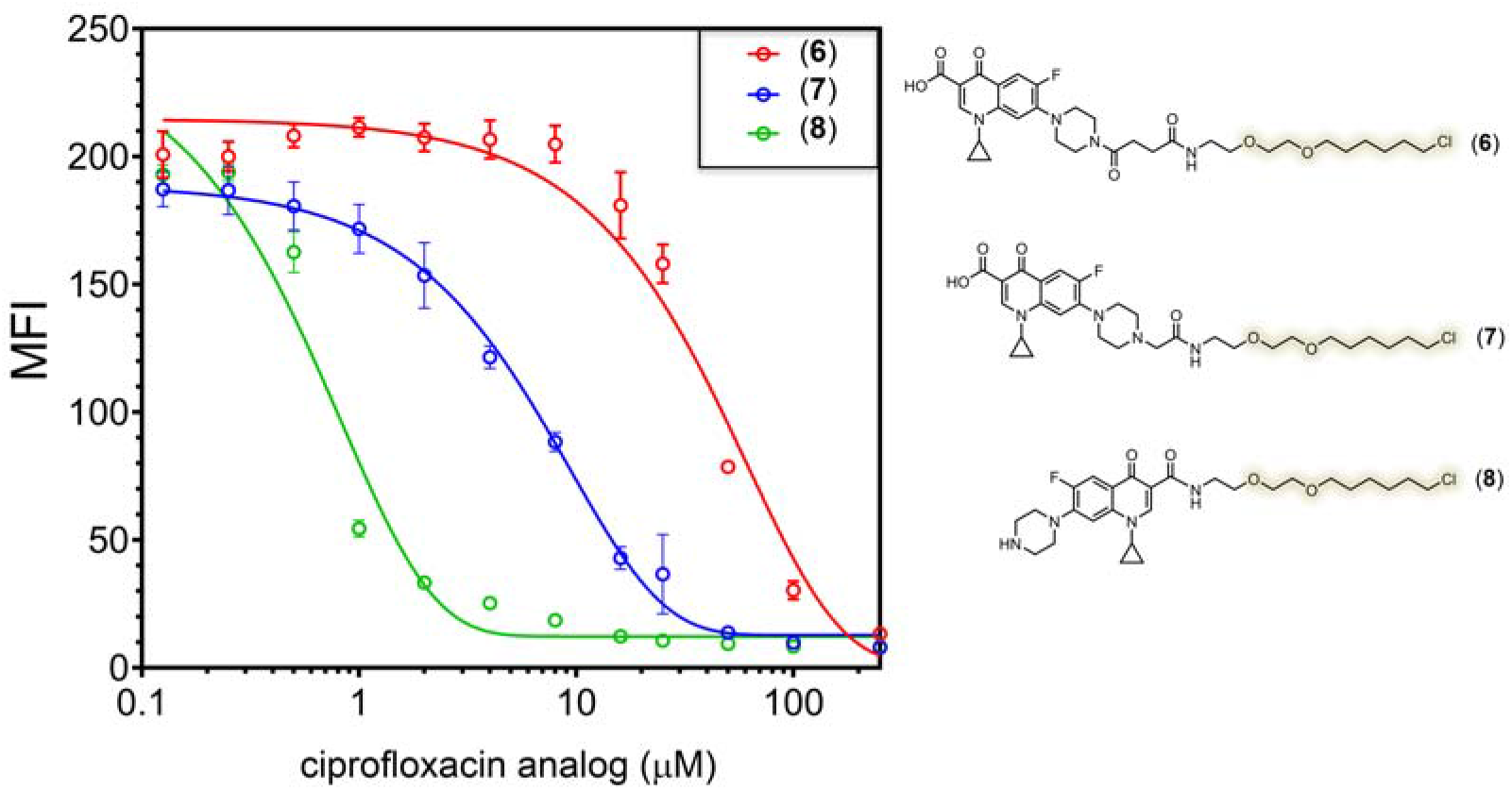
Flow cytometry analysis of *E. coli* treated with various concentrations of molecules **6-8**, followed by a treatment with 5 μM of **R110cl** in the presence and absence of IPTG induction. Data are represented as mean +/− SD (n = 3).

After validating its efficacy in *E.* coli, we optimized the HaloTag system as a reporter for cell envelope permeability in the model mycobacterial species *Mycobacterium smegmatis* and the human pathogen *M. tuberculosis.* We first expressed, in *M. smegmatis* mc^2^155 (NC_008596 in GenBank), HaloTag alone or HaloTag fused to RodA, which encodes the cell wall-synthesizing SEDS family protein^*58–61*^ and subjected HaloTag-expressing *M. smegmatis* to **R110cl**, **TAMRAcl** (**TMRcl**) or **COMcl**. By microscopy and flow cytometry analysis, we found that fluorescent chloroalkane labelling is HaloTag-specific (**Figure 6**). As in *E. coli* (**Figure 2**), **COMcl** labelling was poor (**Figure S8**), so we did not pursue it further. Inspection of the images from **R110cl** and **TMRcl** labelling experiments revealed that the HaloTag alone conferred diffuse, cell-wide fluorescence, consistent with cytoplasmic localization (**Figures 6A,D**), whereas RodA-HaloTag-dependent fluorescence primarily encircled *M. smegmatis* cells, consistent with cell wall localization (**Figures 6C,F**).

**Figure 6.**
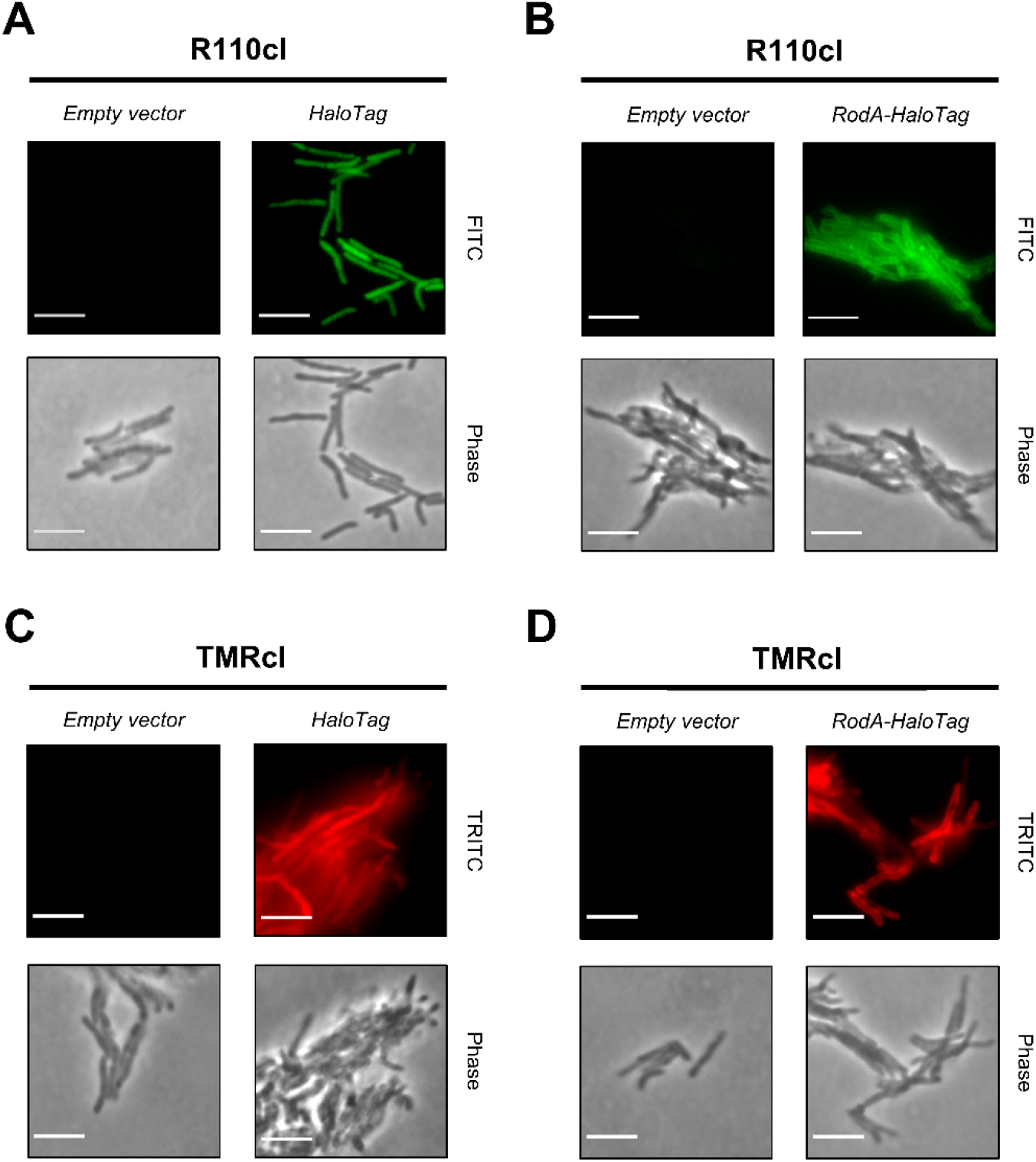
Labelling of HaloTag-expressing *M. smegmatis* strains. Fluorescence microscopy images analysis of *M. smegmatis* transformed with pMV361 HaloTag expressing vector after treatment with **R110cl** (**A**) or **TMRcl** ligand (**C**). The fluorescence distribution suggests HaloTag cytoplasmic localization when expressed as free protein. *M. smegmatis* transformed with pL5ptet0 HaloTag expressing vector and treated with **R110cl** (**B**) or **TMRcl** (**D**). The expression of HaloTag as fusion protein with RodA led to peripheral fluorescence suggesting cell envelope localization. MFI: mean fluorescence intensity. Scale bar 5 μm. Data represents mean and standard deviation of three biological replicates.

The *Corynebacterineae* envelope is distinct from that of other bacteria, making *M. smegmatis* a superior choice for modeling mycobacterial permeability compared to *E. coli,* a phylogenetically distinct, Gram-negative organism. However, there are clear differences in envelope composition between *M. smegmatis* and pathogenic mycobacterial species that have been linked to both virulence and whole cell permeability.^*62*^ Therefore, we expressed HaloTag in *M. tuberculosis* mc^2^6206^*63*^ and subjected HaloTag-expressing *M. tuberculosis* to **R110cl** or **TMRcl**. As in *E. coli* and *M. smegmatis,* we found that fluorescent chloroalkane labelling in *M. tuberculosis* is HaloTag-specific (**Figures 7A-B**). Inspection of the images in **Figure 7B** revealed that HaloTag expression conferred diffuse, cell-wide fluorescence in *M. tuberculosis* cells, again consistent with cytoplasmic localization.

**Figure 7.**
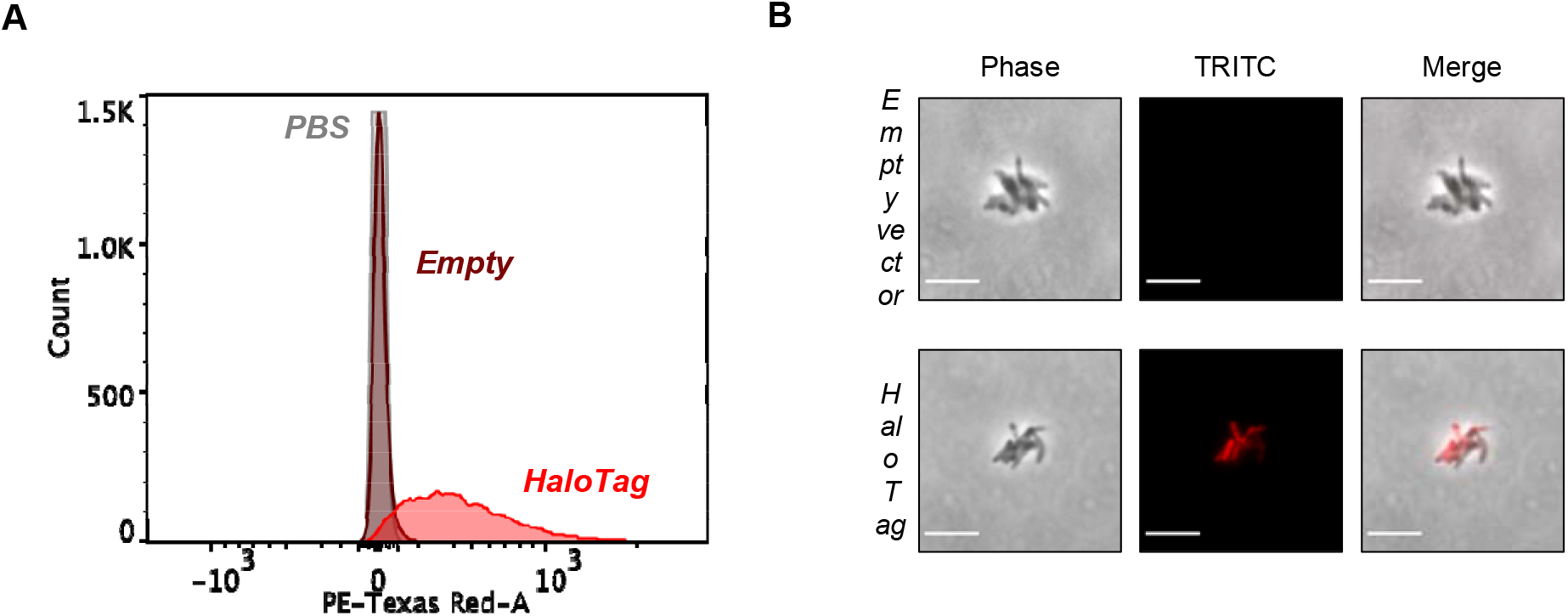
Labelling of HaloTag-expressing *M. tuberculosis.* (**A**) Flow cytometry analysis of *M. tuberculosis* transformed with HaloTag expressing vector and (**B**) relative fluorescence microscope images. Scale bar 5 μm.

Our data suggested that HaloTag could be expressed in both *M. smegmatis* and *M. tuberculosis* and detected by multiple fluorescent chloroalkanes. Using conditions analogous to those that we developed for *E. coli,* we performed a pulse chase experiment in which we pulsed HaloTag-expressing cells with compound (**1**) and chased with **R110cl** (**Figure 8A-C**). For both species we were able to compete **R110cl** fluorescence down to background levels using a 100 μM of compound (**1**) (bacteria transformed with the empty vector, **Figure 8**). We then measured the EC_50_ for (**1**) and (**8**) in *M. tuberculosis* as ~ 5 μM. Similar results were observed when permeability was probed with **TMRcl** (**Figure S9**). These data suggest that HaloTag can quantitatively report the permeation of non-fluorescent chloroalkanes across the mycobacterial cell envelope.

**Figure 8.**
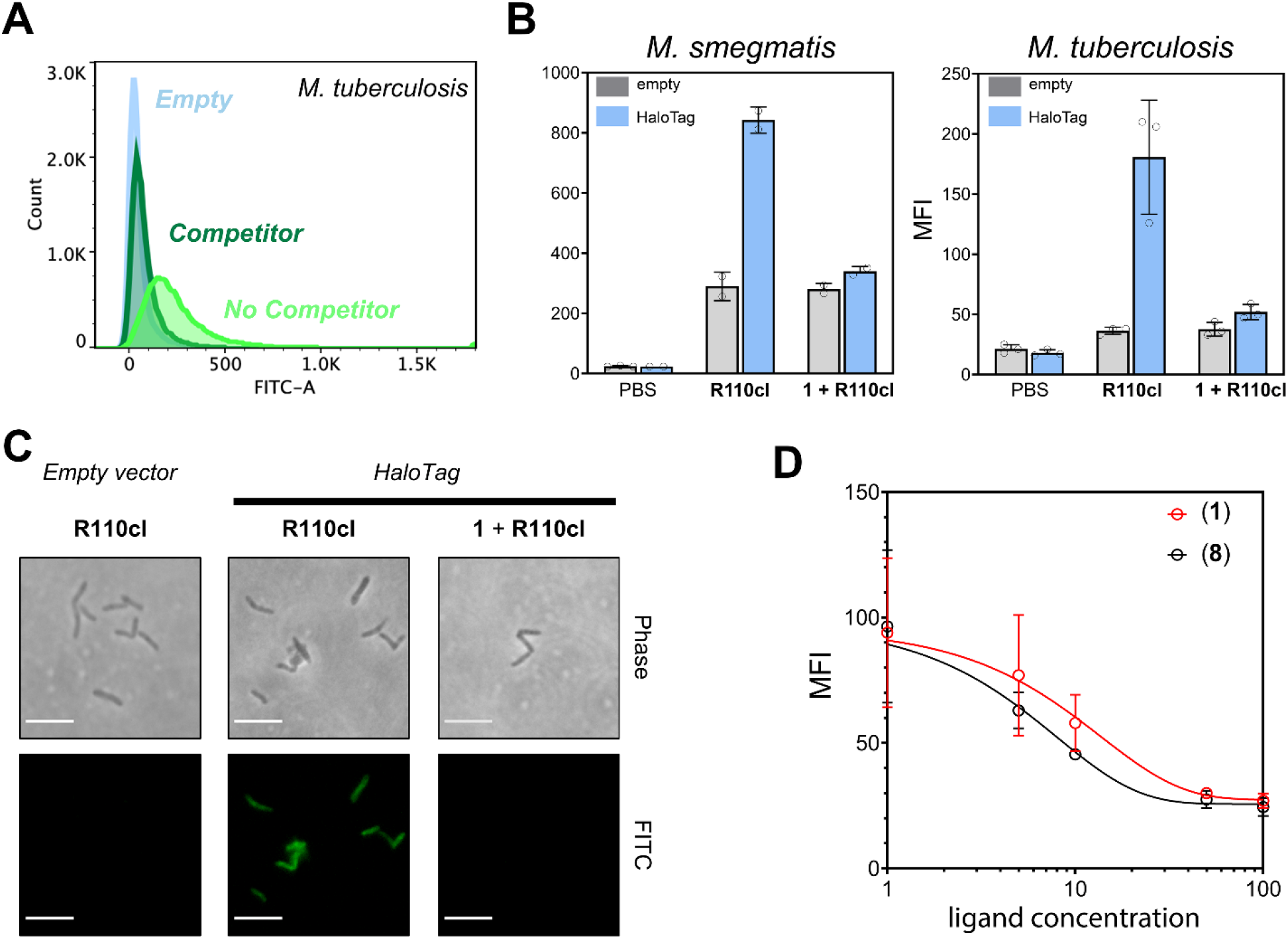
BaCAPA competition assay in *M. smegmatis* and *M. tuberculosis.* (**A**) Flow cytometry analysis of HaloTag-expressing *M. tuberculosis* treated with **R110cl** and pre-treated or not with 100 μM of (**1**). (**B**) Median fluorescence intensity (MFI) of *M. smegmatis (left)* and *M. tuberculosis (right)* of bacteria treated as in **A**. (**C**) Fluorescence microscope images of *M. tuberculosis* underwent BaCAPA competition assay. (**D**) Flow cytometry analysis of *M. tuberculosis* treated with increasing concentrations of the chloroalkane linker (**1**) or ciprofloxacin-chloroalkane (**8**) followed by 1.5 μM **R110cl**. Data represents mean and standard deviation of two (**D**) or three (**B**) biological replicates. Scale bar 5 μm.

Finally, we hypothesized that BaCAPA should also be operational in probing accumulation of small molecules into bacteria within host cells (**Figure 9**). Given that CAPA previously showed low background labeling of mammalian cells in the absence of HaloTag expression, we expected the same when those cells are infected with bacteria.^*40, 64–66*^ The principal reason to establish the feasibility of BaCAPA inside mammalian cells is that a large fraction of serious human pathogens (e.g., *Mycobacterium tuberculosis, Salmonella enterica, Chlamydia trachomatis,* and *Neisseria gonorrhea)* can survive (sometimes exclusively) inside host cells.^*67*^ Moreover, extracellular bacteria can also temporarily reside inside host cells to avoid the action of antibiotics and to promote tissue dissemination.^*68, 69*^ The mammalian plasma membrane can compound the challenges in drug accumulation by introducing an additional barrier. Recent advances in the measurement of accumulation of drugs into bacterial cells have not been extended to bacteria in mammalian cells. To demonstrate a possible iteration of this type of analysis, we set out to show that there was specific labeling of bacterial cells inside macrophages. For *E. coli,* HaloTag protein expression was induced with IPTG prior to exposure to J774 murine macrophage-like cells; after phagocytosis, the co-culture was treated with **R110cl** (**Figure 9A**). Our results revealed that there was a large increase in fluorescence levels when J774 cells were incubated with HaloTag expressing bacterial cells (**Figure 9B**). The absence of HaloTag induction and J774 cells alone showed background levels of fluorescence. The same bacterial cells were also subjected to treatment with **R110cl** (with no host cell incubation) and were found to have, as expected, high levels of fluorescence when induced with IPTG (**Figure S10**). Confocal microscopy confirmed that cellular fluorescence was exclusively localized to bacteria (**Figure 9C**).

**Figure 9.**
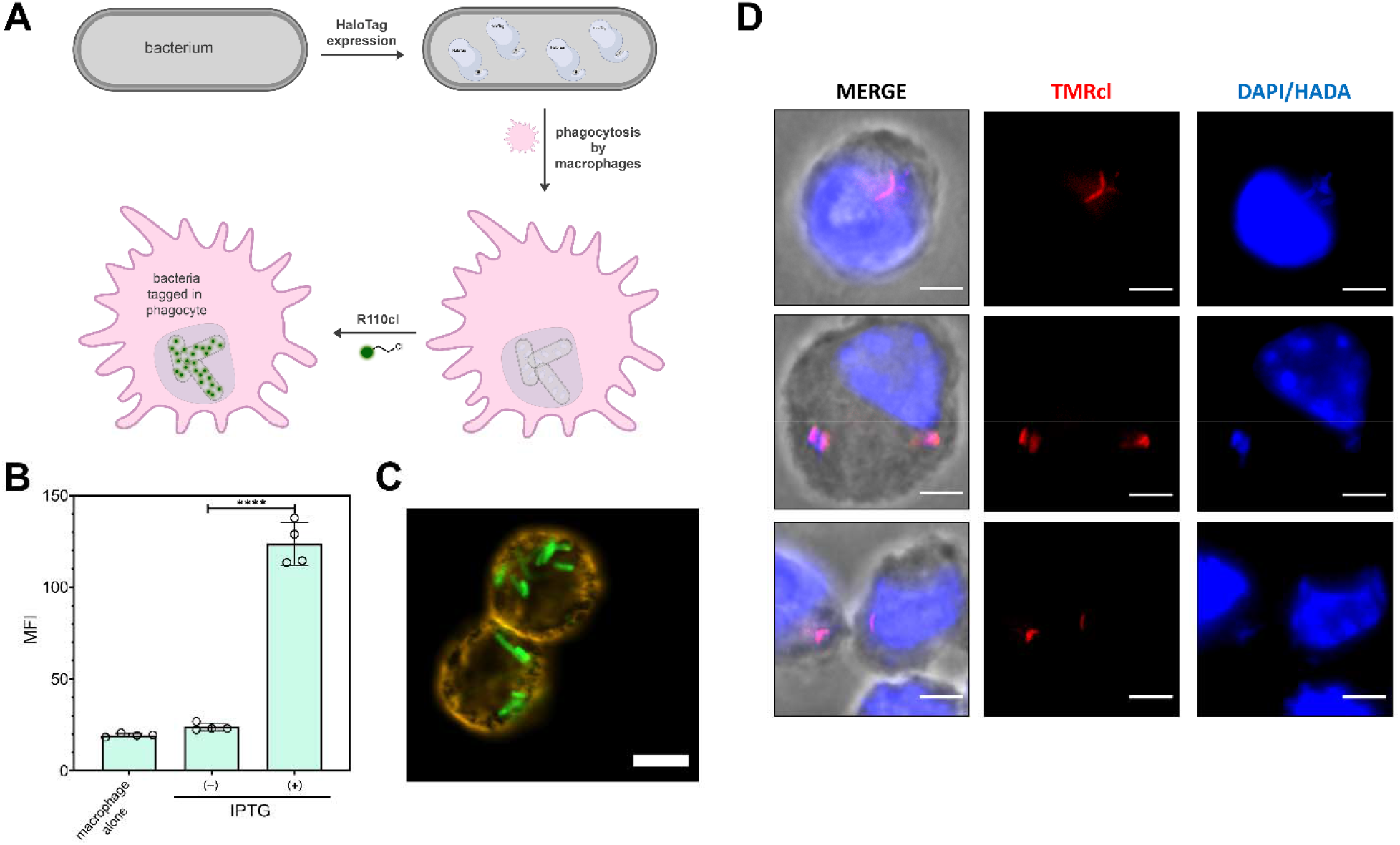
(A) Schematic representation of BaCAPA of bacteria inside mammalian cells. (B) Flow cytometry analysis of J774 cells alone, or J774 cells phagocytosed *E. coli* in the presence and absence of IPTG induction treated with 1.5 μM of **R110cl**. Data are represented as mean +/− SD (n = 3). *P*-values were determined by a two-tailed *t*-test (* denotes a *p*-value < 0.05, **□<□0.01, ***<0.001, ns□=□not significant). (C) Confocal microscopy images of J774 macrophages with phagocytosed *E. coli* that were treated with 5 μM of **R110cl**. Cells were fixed with formaldehyde and treated with tetramethyl-rhodamine-tagged WTA (5 μg/mL) for 30 min. Shown is the overlay of the channels corresponding to rhodamine 110 and tetramethyl-rhodamine. Scale bar = 10 μm. (D) Fluorescent microscopy of iBMDM infected with HaloTag-expressing *M. tuberculosis* after treatment with 1 μM. **TMRcl**. Scale bar 5 μm.

A phagosome occupied by a non-pathogenic microbe, *e.g., E. coli* K12, has different characteristics than that manipulated by a pathogen such as *M. tuberculosis.* Moreover, the bacteria themselves respond differently (death vs. survival/growth) to the immune insults of a professional phagocytic cell like a macrophage. Accordingly, we tested whether we could detect HaloTag labeling in *M. tuberculosis* within immortalized C57BL/6 mouse bone marrow derived macrophages (iBMDM). *M. tuberculosis* transformed with empty or HaloTag-expressing pMV361 vector were pre-labeled with the cell wall-incorporating blue fluorescent probe HADA^*70–73*^, washed, then used to infect iBMDM. We were then able to visualize HADA-labeled HaloTag-expressing *M. tuberculosis* (but not the negative control strain) after a brief incubation of the co-culture in **TMRcl** (**Figure 9D**). Taken together, our results establish the ability of BaCAPA to detect phylogenetically-distinct diderms *E. coli* and *M. tuberculosis* within host cells.

In conclusion, we have shown that BaCAPA is able to readily report on the accumulation of small molecules in *E. coli, M. smegmatis* and *M. tuberculosis via* irreversible covalent bond formation with HaloTag. The expression of HaloTag in the cytoplasm or periplasm of *E. coli* and *M. smegmatis* resulted in labeling that was consistent with the expected localization of the protein. Fluorescence signals with rhodamine-based (and, in mycobacteria, TAMRA-based) dyes proved to be superior to coumarin, thus resulting in sensitive measurements. We first demonstrated that BaCAPA can also be adopted to establish the disruption of the outer membrane of *E. coli* as demonstrated by PMBN and liproxstatin. Next, we assembled a library of small molecules containing an amino group with varying levels of methyl modifications and found that for this group of molecules, the methylation level was mostly independent of the apparent accumulation levels in *E. coli* cells. We also synthesized a group of ciprofloxacin analogs and observed that there was a wide range of apparent accumulation based on the structural modifications. Most significantly, there was a prominent discord between apparent accumulation levels and the MIC values of these modified ciprofloxacin molecules in *E. coli.* These results highlight the potential challenges in using MIC values as a proxy for drug accumulation, which can confound assignments for structure-activity relationships. We also demonstrated the ability of BaCAPA to quantitatively report permeation across *M. tuberculosis* cell envelope. Finally, we showed that BaCAPA is compatible with both non-pathogenic *E. coli* and pathogenic *M. tuberculosis* within mammalian cells. Together with our pulse chase data these experiments suggest the potential for BaCAPA to detect drug accumulation across multiple host and bacterial membranes. Our results establish BaCAPA as a robust and readily adoptable fluorescence-based reporter for compartment-specific accumulation of molecules into bacterial cells.

## SUPPORTING INFORMATION

Additional experimental details (methods, characterization and synthesis of peptide probes) and figures. This material is available free of charge via the Internet at http://pubs.acs.org.

## ACKNOWLEDGEMENT

This study was supported by the NIH grant GM124893-01 (M.M.P.) and DP2 AI138238 (M.S.S.). We acknowledge the Keck Center for Cellular Imaging for the usage of the Zeiss 980 microscopy system (PI:AP; NIH-OD025156). We would like to thank Dr. Junhao Zhu for helpful discussion and sharing of *M. tuberculosis* labeling protocols and Dr. Amy Burnside, Facility Director of the University of Massachusetts Amherst Flow Cytometry Facility at the Institute for Applied Life Sciences.

## TABLE OF CONTENTS GRAPHIC

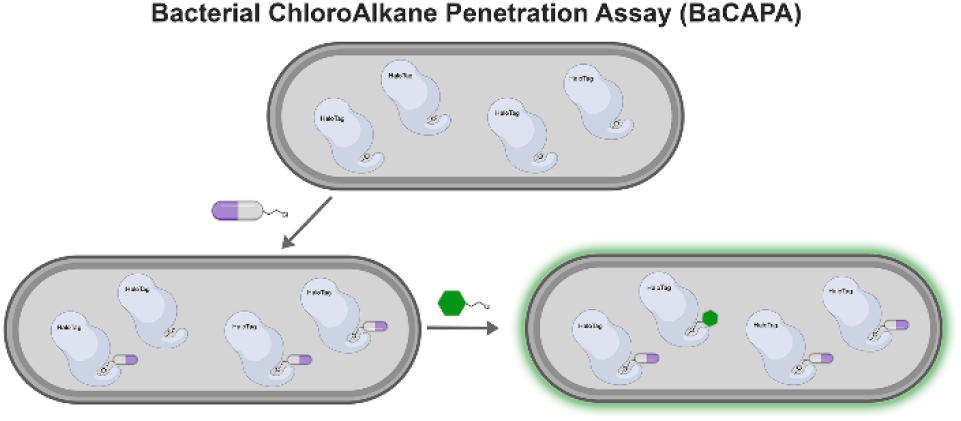

