## Supporting Information for "Compartment-Specific Measurement of Small Molecule Accumulation into Diderm Bacteria"

#### Supplementary Figures

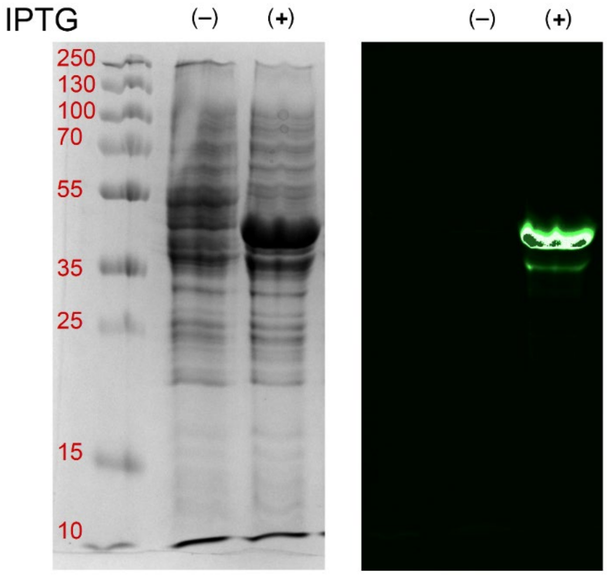


**Figure S1.** Coomassie (left) and fluorescence images of SDS-PAGE gel of whole cellular extracts of *E. coli* treated with 5 μM of **R110cl** in the presence and absence of IPTG induction.

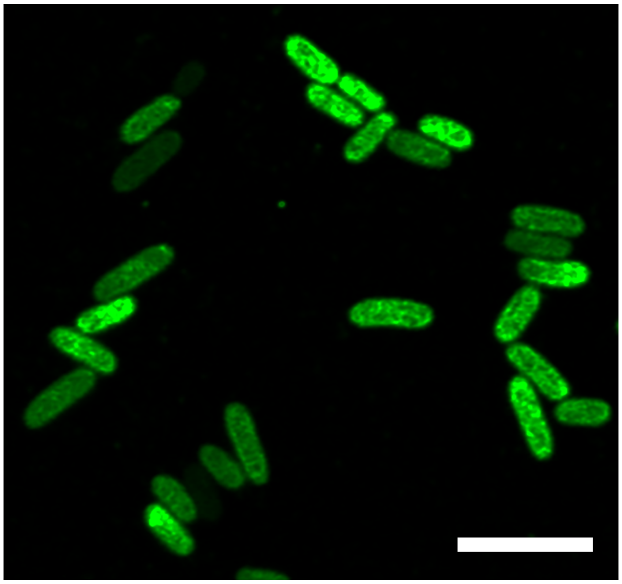

**Figure S2.** Confocal microscopy analysis of *E. coli* BL21(DE3) expressing cytoplasmic HaloTag induced with IPTG and treated with 5 μM of **R110cl**. Cells were fixed with 2 % formaldehyde, deposited onto a pre-cooled 1% (w/v) agarose pad placed on a glass microscope slide, and covered with a micro cover glass. Images were collected with a Zeiss 880/990 multiphoton Airyscan microscopy system (60x oil-immersion lens). Scale bar = 2 μm.

| | | |
|---|---|---|
| 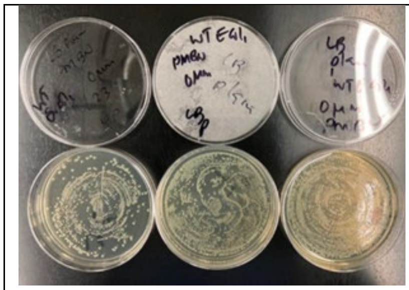 0 μM PMBN: >300 colonies | 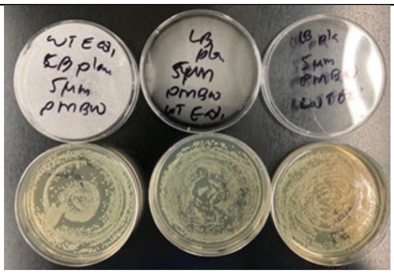 5 μM PMBN: >300 colonies | 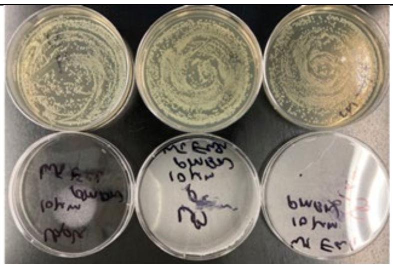 10 μM PMBN: >300 colonies |
| 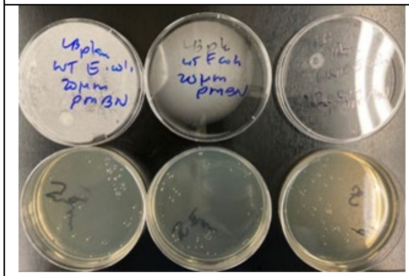 20 μM PMBN: 51 colonies | 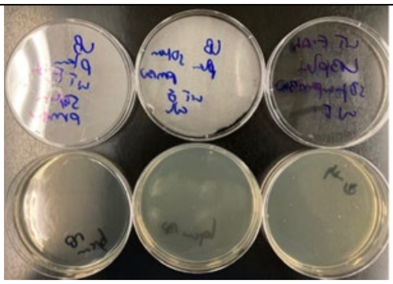 50 μM PMBN: 0 colonies | 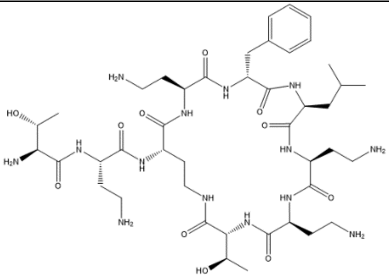 PMBN, polymyxin nonapeptide |

**Figure S3.** *E. coli* BL21(DE3) cells were treated with stated concentrations of PMBN.

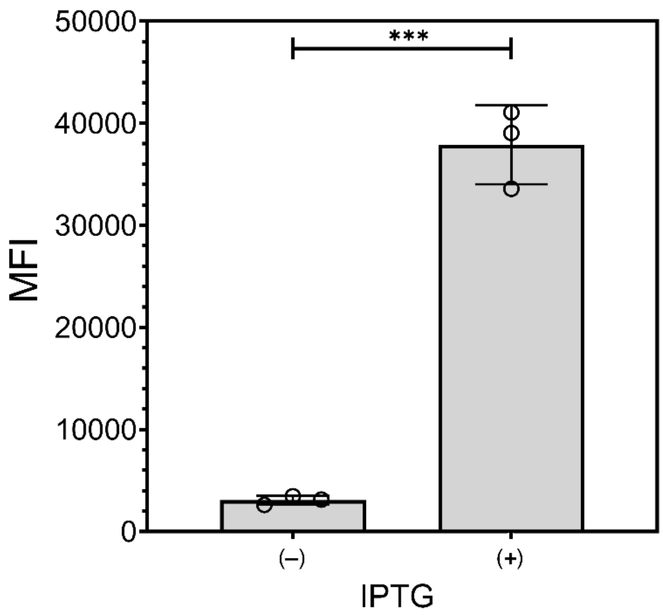


**Figure S4.** *E. coli* was treated with 5 μM of **R110cl** in the presence and absence of IPTG induction. After washing cells with PBS (1X), cellular fluorescence was measured using a plate reader (excitation = 480 nm (40 nm), emission = 520 nm (40 nm)).

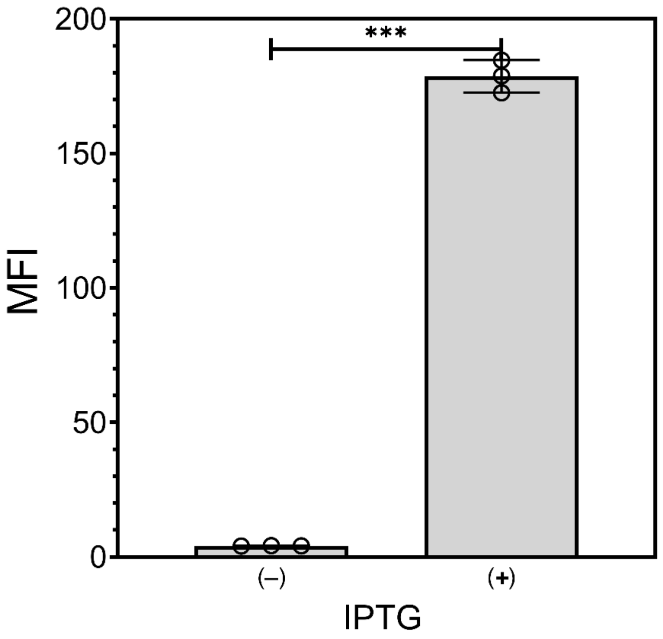


**Figure S5.** Flow cytometry analysis of *E. coli* (ATCC 25922) expressing cytoplasmic HaloTag treated with 5 μM of **R110cl** in the presence and absence of IPTG induction. Data are represented as mean +/- SD (n = 3). *P*-values were determined by a two-tailed *t*-test (* denotes a *p*-value < 0.05, ** < 0.01, ***<0.001, ns = not significant).

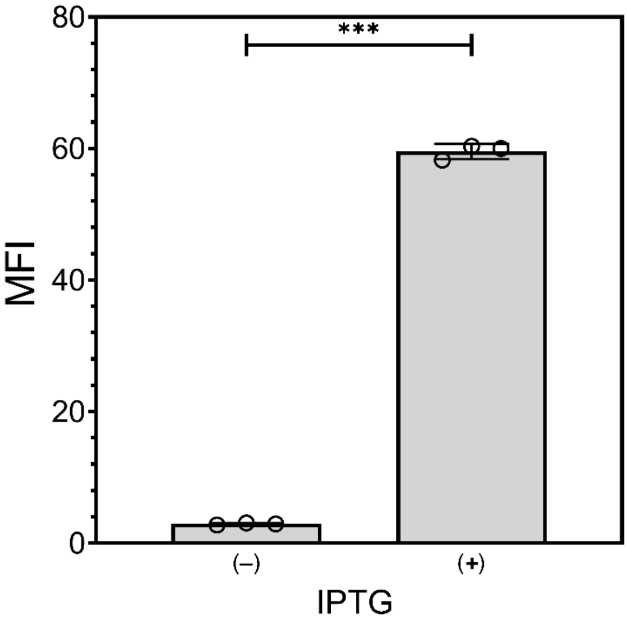


**Figure S6.** Flow cytometry analysis of *E. coli* (Lemo) carrying a plasmid of HaloTag fused with DsbA signal peptide treated with 5 μM of **R110cl** in the presence and absence of IPTG induction. Data are represented as mean +/- SD (n = 3). *P*-values were determined by a two-tailed *t*-test (* denotes a *p*-value < 0.05, ** < 0.01, ***<0.001, ns = not significant).

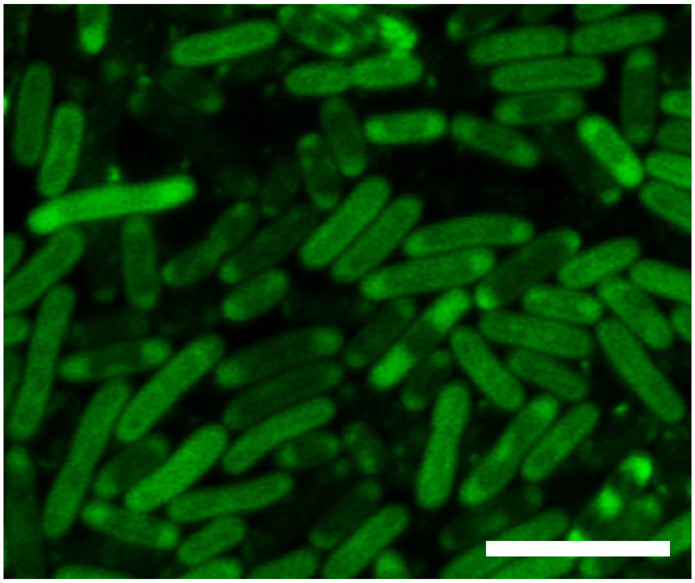

**Figure S7.** Confocal microscopy analysis of *E. coli* (Lemo) carrying a plasmid of HaloTag fused with DsbA signal sequence treated with 5 μM of **R110cl** in the presence of IPTG induction. Cells were fixed with 2 % formaldehyde, deposited onto a pre-cooled 1% (w/v) agarose pad placed on a glass microscope slide, and covered with a micro cover glass. Images were collected with a Zeiss 880/990 multiphoton Airyscan microscopy system (60x oil-immersion lens). Scale bar = 2μm.

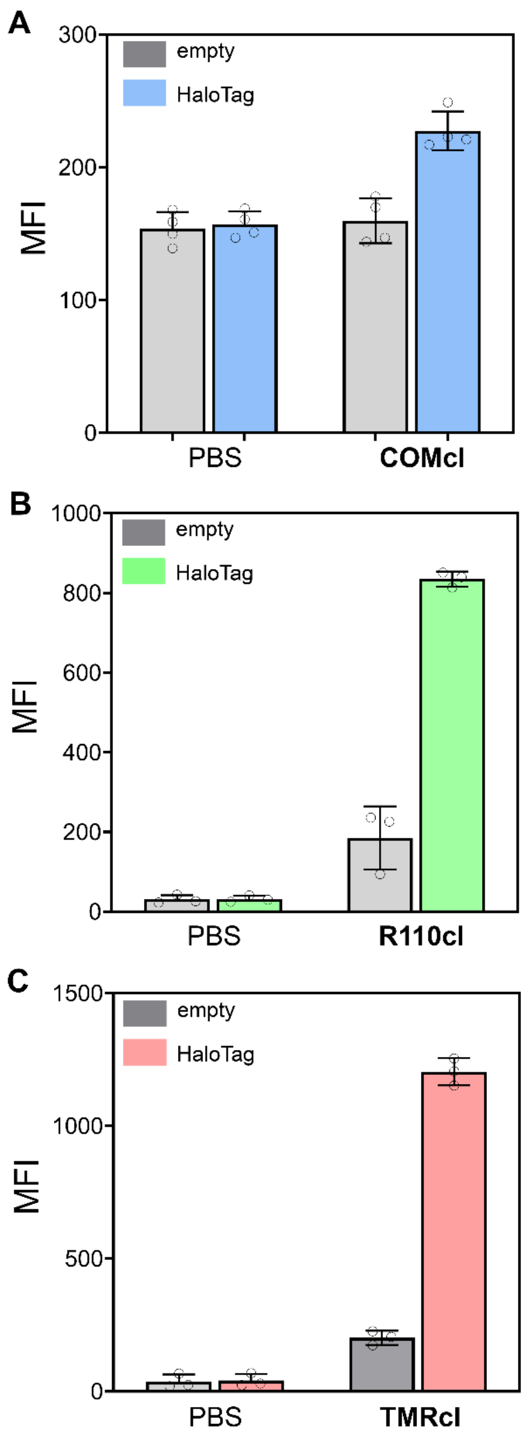


**Figure S8. Labelling of HaloTag-expressing *M. smegmatis* with various dyes.** Flow cytometry analysis of *M. smegmatis* transformed with *pMV361_HaloTag* vector after treatment with (A) **COMcl**, (B) **R110cl**, and (C) **TMRcl**. MFI: median fluorescence intensity. Data represents mean and standard deviation of three biological replicates.

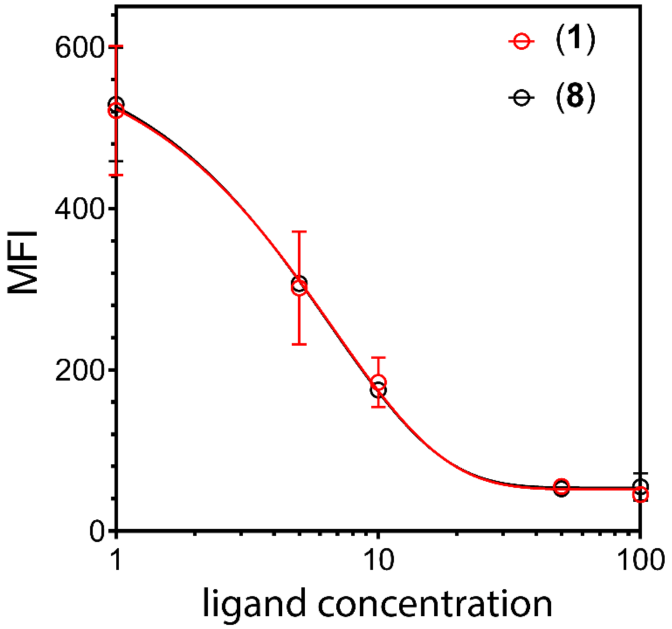


**Figure S10.** Flow cytometry analysis of *M. tuberculosis* treated with increasing concentrations of the chloroalkane linker (**1**) or ciprofloxacin-chloroalkane (**8**) followed by 0.25 μM **TMRcl**.

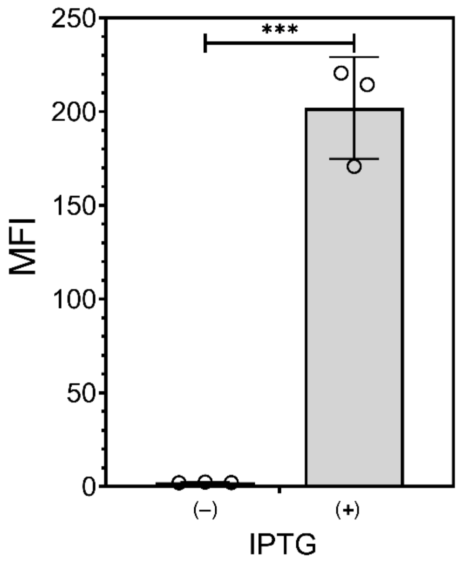


**Figure S10.** Flow cytometry analysis of *E. coli* treated with 5 μM of **R110cl** in the presence and absence of IPTG induction. The same cells were used for the analysis of labeling bacteria in macrophages. Data are represented as mean +/- SD (n = 3). *P*-values were determined by a two-tailed *t*-test (* denotes a *p*-value < 0.05, ** < 0.01, ***<0.001, ns = not significant).

**Materials & Methods**

**General**

All reagent grade reagents were purchased from commercial vendors and were used without further purification. Anhydrous solvents dichloromethane (DCM), methanol, tetrahydrofuran (THF), and dimethylformamide (DMF) were purchased from Sigma. LB Broth (Miller formulation), Mueller Hinton Broth 2, triethylamine, bromoacetyl bromide, and ciprofloxacin were also purchased from Sigma. Vancomycin hydrochloride, anhydrous betaine, Boc-sarcosine N-hydroxysuccinimide ester, and Boc-glycine N-hydroxysuccinimide ester were all purchased from ChemImpex. N,N-Dimethylglycine and di-isopropyl ethylamine (DIPEA) were both purchased from TCI America. 2-(2-((6-Chlorohexyl)oxy)ethoxy)ethanamine hydrochloride was purchased from AmBeed; this compound is called HaloTag amine ligand hereinafter. HaloTag® Coumarin Ligand (chloroalkane-tagged coumarin) and HaloTag® R110Direct™ Ligand (chloroalkane-tagged rhodamine 110) were both purchased from Promega. All reaction vessels were flame dried prior to use. Reaction progress with ciprofloxacin was monitored by thin-layer chromatography, visualized with UV light. Compounds were purified via reversed phase HPLC using Luna 10μm C8(2) 100 Å LC Column 250 x 21.2mm (Phenomenex) with a Waters 1525 Binary HPLC solvent pump. Purity of compounds was confirmed via reversed phase HPLC using a Luna 5μm C8(2) 100 Å LC Column 250 x 4.6mm (Phenomenex) with a Waters 1525 Binary HPLC solvent pump. UV-visible spectra were collected on Thermo Scientific Genesys-50 spectrophotometer using either transparent plastic or quartz cuvettes. $^{1}H$ and $^{13}C$-NMR spectra for all new compounds and intermediates for characterization were acquired on a Varian 600MHz spectrophotometer. All NMR spectra were analyzed using MestreNova software. Residual solvent signal from $CDCl_3$, $CD_3OD$ and DMSO-d6 referenced to tetramethylsilane (TMS) were used as reference standards for defining chemical shifts of $^1H$ or $^{13}C$ spectra of compounds. Chemical shifts are reported in $\delta$ ppm and coupling constants (J) are reported in Hertz [Hz]. Deuterated solvents were used as received from Cambridge Isotopes. Routine mass analysis was performed on Advion Expression® CMS mass spectrometer using standard parameters for intermediates. For the analysis of fragmentation sensitive compounds, low fragmentation, low energy setup was used. The observed molecular weights for compounds were represented as m/z. High resolution electrospray ionization mass spectrometry (HRMS, ESI/MS) analyses were obtained on an Agilent 6545B Q-TOF LC/MS equipped with 1260 Infinity II LC system with auto sampler. Samples were dissolved in either methanol or MeCN and eluted with a MeCN/$H_2O$ solution containing 0.1% formic acid. Fluorescence measurement for the probes was performed on Synergy H1 multimode hybrid microplate reader from BioTek with appropriate setting for fluorophores. The HPLC fractions of the desired purified compounds were first concentrated under reduced pressure using rotary evaporator Hei-CHILL (Heidolph). The final concentrated aqueous solutions were lyophilized to dryness using Labconco Freezone 4.5L (-84$^{o}$C) lyophilizer.

### Synthesis and characterization of chloroalkane-tagged derivatives

#### Synthesis of glycinamide-HaloTag derivatives:

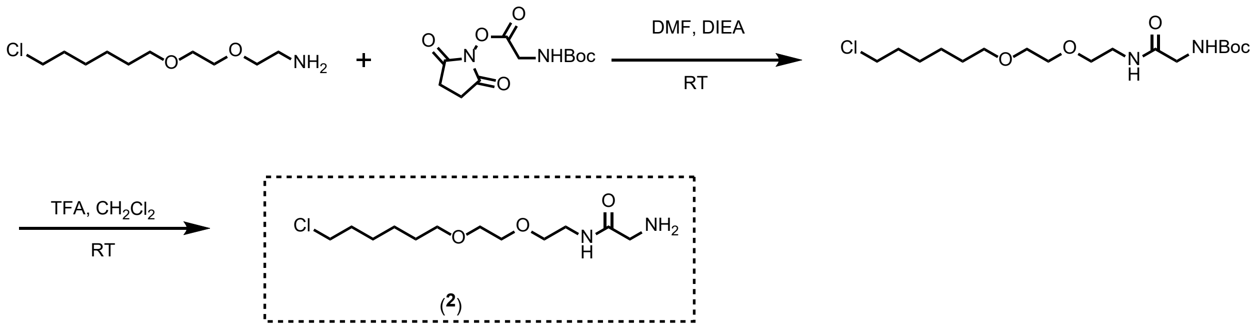


Compound (**2**), HaloTag-glycinamide (R = H): To a solution of N-O$^t$Boc-glycine-NHS ester (55mg, 0.2 mmol) in DMF (3.0mL) was first added DIEA (300μL), followed by HaloTag amine ligand (52.0mg, 0.2mmol) at room temperature. The mixture was stirred overnight, diluted with ether (20 mL), washed with dil. HCl (0.1M), water and finally with 10% $NaHCO_3$ solution. The product was dried over $Na_2SO_4$, concentrated under reduced pressure, and the leftover product was used as is. The N- -$^t$Boc deprotection was performed with 10% TFA in $CH_2Cl_2$. The mixture was concentrated and subjected to preparative HPLC purification. The fractions showing the desired mass were collected, concentrated under reduced pressure, and finally lyophilized to yield a transparent syrup (39.0mg, yield: 69%). $^1$H-NMR ($CDCl_3$)δ1.34 (m, 2H, $CH_2$), 1.44 (m, 2H, $CH_2$), 1.57 (m, 2H, $CH_2$), 1.76 (m, 2H, $CH_2$), 3.44-3.48. (m, 4H, $CH_2$), 3.50-3.60 (m, 8H, $OCH_2$), 3.78 (m, 2H, $CH_2$), 7.76 (brs, 1H, NH), 8.11 (brs, 2H, $NH_2$). $^{13}$C-NMR ($CDCl_3$) δ 25.4, 26.8, 29.3, 32.6, 39.7, 41.2, 45.2, 69.5, 69.8, 70.2, 71.3, 166.7. HRMS: Obs. 281.1632 for [M+H]; Calc. for $C_{12}H_{25}ClN_2O_3$ 280.1554.

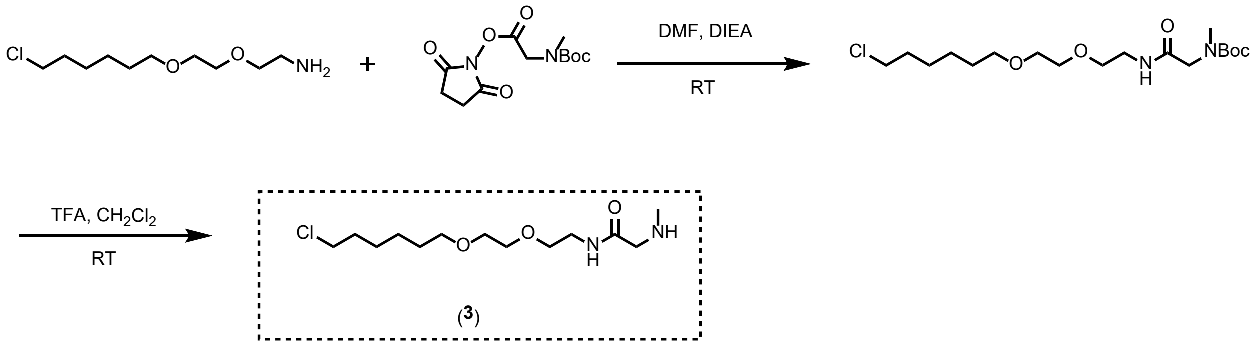


Compound (**3**), HaloTag-N-methyl glycinamide: Following the foregoing procedure, except for the use of N-$^{t}$Boc-sarcosine-NHS ester (61.0mg, 0.21mmol), afforded a transparent syrup (47.0mg, yield: 76%). $^{1}$H-NMR ($CDCl_3$)δ1.35 (m, 2H, $CH_2$), 1.44 (m, 2H, $CH_2$), 1.59 (m, 2H, $CH_2$), 1.77 (m, 2H, $CH_2$), 2.80 (brs, 3H, $NCH_3$), 3.47 (m, 4H, $CH_2$), 3.53 (t, *J* = 6 Hz 4H, 2x $CH_2$), 3.54-3.59 (m, 10H, $CH_2$), 3. 85 (brs, 2H, $CH_2$), 7.53 (brs, 1H, NH), 8.10 (brs, 1H, NH). $^{13}$C-NMR ($CDCl_3$) δ 25.4, 26.7, 29.3, 32.6, 33.9, 39.8, 45.1, 50.5, 69.3, 69.9, 70.3, 71.4, 165.3. HRMS: Obs. 295.1806 for [M+H]; Calc. for $C_{13}H_{27}ClN_2O_3$ 294.1710.

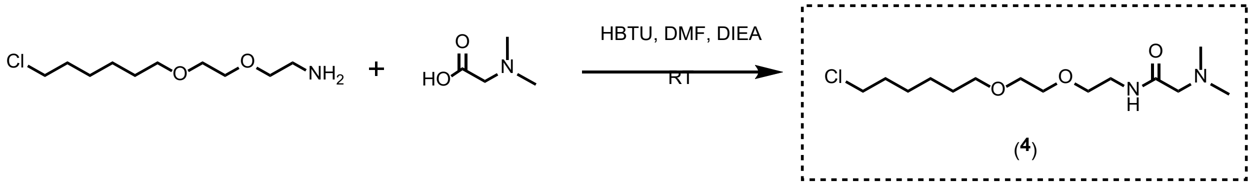


Compound (**4**), HaloTag-N, N-Dimethyl glycinamide: To a solution of N, N-dimethylglycine (33 mg, 0.32mmol) in anhydrous DMF (2.0 mL), DIEA (250μL, 1.43 mol) and HBTU (185 mg, 0.49mmol) were added and stirred for 10 minutes. Subsequently, HaloTag amine ligand (86mg, 0.335mmol) was added to the mixture at room temperature. The mixture was stirred overnight, diluted with ether (25 mL), washed with water and 10% $NaHCO_3$ solution, dried over $Na_2SO_4$, and concentrated under reduced pressure. The crude product was purified by preparative HPLC. The fractions showing m/z 309 [M+H] were collected, concentrated under reduced pressure, and finally lyophilized to yield a transparent syrup (71.0mg, yield: 72%). $^1$H-NMR ($CDCl_3$)δ1.34 (m, 2H, $CH_2$), 1.45 (m, 2H, $CH_2$), 1.58(m, 2H, $CH_2$), 1.78 (m, 2H, $CH_2$), 2.97 (s, 6H, $NCH_3$), 3.44-3.48 (m, 4H, $CH_2$), 3.52-3.59 (m, 8H, 4 x $CH_2$, $OCH_2$), 3.83 (s, 2H, $CH_2$), 7.99 (brs, 1H, NH). $^{13}$C-NMR ($CDCl_3$) δ 25.4, 26.7, 29.5, 32.6, 38.7, 45.0, 46.0, 63.16, 70.0, 70.1, 70.3, 71.3, 170.7. HRMS: Obs. 309.1970 for [M+H]; Calc. for $C_{14}H_{30}ClN_2O_3$ 309.1939.

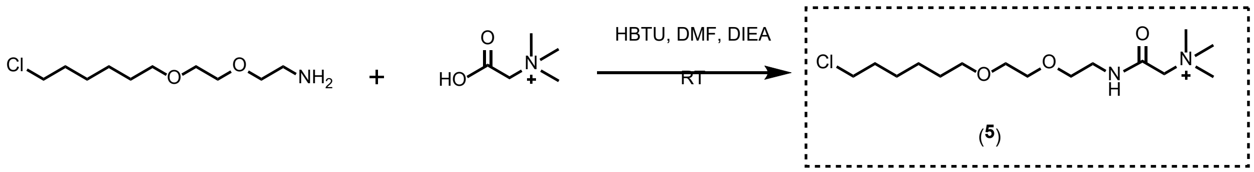


Compound (**5**), HaloTag-betainamide: To a solution of betaine (35.5mg, 0.3mmol) in 1mL DMF was added DIEA (250μL, 1.43 mmol) and HBTU (185 mg, 0.49mmol), and stirred for 10 minutes. Subsequently, HaloTag amine ligand (86mg, 0.335mmol) was added to the mixture at room temperature. The mixture was stirred overnight, diluted with ether (25 mL), washed with water and 10% $NaHCO_3$ solution, dried over $Na_2SO_4$, and concentrated under reduced pressure. Upon reaction, the mixture was diluted with HPLC solvent (9 mL, 35% ACN: 65% water, with 0.1%TFA) and was directly loaded on preparative HPLC. Using the same gradient used for other compounds, fractions showing m/z 323 $[M]^+$ were collected, concentrated under reduced pressure, and finally lyophilized to yield a transparent syrup (57 mg, yield: 59%). $^1$H-NMR ($CDCl_3$)δ1.35 (m, 2H, $CH_2$), 1.45 (m, 2H, $CH_2$), 1.57 (m, 2H, $CH_2$), 1.76 (m, 2H, $CH_2$), 3.35 (s, 9H, $CH_3$), 3.44 (t, 2H, $CH_2$), 3.47 (m, 2H, $CH_2$), 3.50-3.60 (m, 8H, $OCH_2$), 4.28 (s, 2H, CH2), 8.39 (brs, 1H, NH). $^{13}$C-NMR ($CDCl_3$) δ 25.4, 26.7, 29.6, 32.6, 39.5, 45.2, 54.7, 65.4, 68.9, 70.0, 70.3, 71.3, 162.8. HRMS: Obs. 323.2176 for $[M+H]^+$ Calc. for $C_{15}H_{32}ClN_2O_3$ 321.2096.

Synthesis of compound **(6)**:

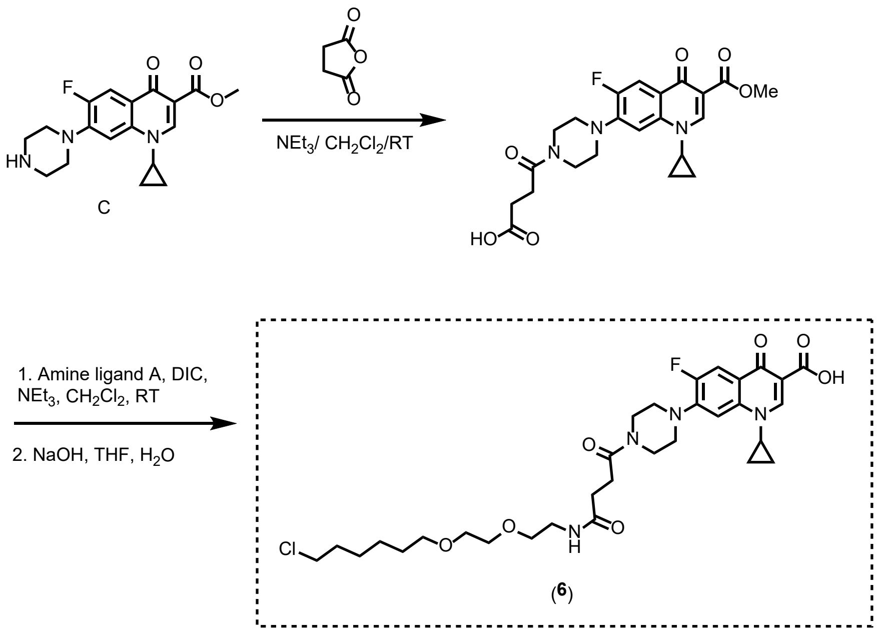


Compound (**6**). Synthesis of N-succinamide of ciprofloxacin methyl ester: The Ciprofloxacin methyl ester as described below (86 mg, 0.25 mmol) was dissolved in 5 mL of anhydrous $CH_2Cl_2$ and to this, $NEt_3$ (100 mg, 1 mmol) was added. To this mixture, succinic anhydride was added as solid (30 mg, 0.3 mmol) and the homogenous mixture stirred at room temperature overnight. TLC analysis revealed that the starting methyl ester was consumed to form a new, non-polar compound. The reaction mixture was diluted with $CH_2Cl_2$ (20mL), transferred to a separatory funnel, and to this, dil. HCl (5 mL, 0.1M) was added. The aqueous layer was separated, and the organic layer was further washed with water and brine. The organic layer was dried over $Na_2SO_4$ and concentrated under reduced pressure. Column chromatography purification over silica-gel (hexanes/ethyl acetate, 1:1) afforded a pure white solid (82mg, yield: 74%). $^1$H-NMR: ($CDCl_3$)$\delta$1.14 (m, 2H, cyclopropyl-$CH_2$), 1.32 (m, 2H, cyclopropyl-$CH_2$), 1.39-1.49 (m, 6H,

3 x $CH_2$), 1.49 (s, 9H, 3 x$CH_3$), 1.58 (dt, *J* = 6Hz, 128Hz, 2H, $CH_2$), 1.74 (dt, *J* = 6, 12Hz, 2H, $CH_2$), 3.22 (t, *J* = 6Hz, 4H, 2 x $CH_2$), 3.43 (m, 1H, CH), 3.45-3.51 (m, 4H, piperidyl 2 x CH2), 3.62 (m, 2H), 3.63-3.67 (m, 4H, 2 x $CH_2$), 7.31 (d, *J* = 6 Hz, 1H, ArH), 8.03 (d, *J* = 12 Hz, 1H, ArH), 8.81 (s, 1H, ArH), 10.1 (brs, 1H, NH). $^{13}$C-NMR: ($CDCl_3$) $\delta$ 8.2, 25.4, 26.8, 28.4, 29.5, 32.6, 34.7, 39.2, 45.1, 50.0, 70.0, 70.2, 70.6, 71.3, 105.0, 111.4, 112.7, 112.8, 122.2, 138.4, 144.8, 146.8, 152.6, 154.6, 165.1, 175.4.

**Synthesis of HaloTag amide from N-succinyl ciprofloxacin methyl ester:** N-Succinyl-ciprofloxacin methyl ester (45mg, 0.1mmol) was dissolved in $CH_2Cl_2$ containing $NEt_3$ (101mg, 1 mmol). To this, DIC (20μL, 16.1 mg, 0.13 mmol) and HaloTag amine ligand (31 mg, 0.12mmol) were added and stirred at room temperature overnight. TLC analysis showed that the starting material was completely consumed to form a new compound of a different polarity. The reaction mixture was concentrated and purified by silica-gel column chromatography with $CHCl_3$ / MeOH (98:2) eluent. The pure compound was isolated after concentration and drying (51mg, yield: 78%). $^1$H-NMR: ($CDCl_3$)$\delta$1.14 (m, 2H, cyclopropyl-$CH_2$), 1.32 (m, 2H, cyclopropyl-$CH_2$), 1.39-1.49 (m, 6H, 3 x $CH_2$), 1.49 (s, 9H, 3 x$CH_3$), 1.58 (dt, *J* = 6Hz, 128Hz, 2H, $CH_2$), 1.74 (dt, *J* = 6, 12Hz, 2H, $CH_2$), 3.22 (t, *J* = 6Hz, 4H, 2 x $CH_2$), 3.43 (m, 1H, CH), 3.45-3.51 (m, 4H, piperidyl 2 x CH2), 3.62 (m, 2H), 3.63-3.67 (m, 4H, 2 x $CH_2$), 7.31 (d, *J* = 6 Hz, 1H, ArH), 8.03 (d, *J* = 12 Hz, 1H, ArH), 8.81 (s, 1H, ArH), 10.1 (brs, 1H, NH). $^{13}$C-NMR: ($CDCl_3$) $\delta$ 8.2, 25.4, 26.8, 28.4, 29.5, 32.6, 34.7, 39.2, 45.1, 50.0, 70.0, 70.2, 70.6, 71.3, 105.0, 111.4, 112.7, 112.8, 122.2, 138.4, 144.8, 146.8, 152.6, 154.6, 165.1, 175.4.

**Hydrolysis of the methyl ester:** The ciprofloxacin-N-acylated HaloTag amide (33 mg, 0.05mmmol) was dissolved in THF (2.0mL) and to this was added 1.0M NaOH (200μL). The mixture was vigorously stirred at room temperature for 12 hrs. TLC analysis showed that the methyl ester was consumed to form a new polar compound. The reaction mixture was concentrated under reduced pressure and acidified with dil. HCl (1M) and extracted with methylene chloride. The organic layer was washed, dried over $Na_2SO_4$, and evaporated under reduced pressure to yield the crude residue which was purified with silica gel column chromatography $CHCl_3$/MeOH (98:2). The pure compound (Compound **(6)**) was isolated after concentration and drying of fractions as a white solid (51mg, yield: 78%). $^1$H-NMR: ($CDCl_3$)δ1.14 (m, 2H, cyclopropyl-$CH_2$), 1.32 (m, 2H, cyclopropyl-$CH_2$), 1.39-1.49 (m, 6H, 3 x $CH_2$), 1.49 (s, 9H, 3 x$CH_3$), 1.58 (dt, *J* = 6Hz, 128Hz, 2H, $CH_2$), 1.74 (dt, *J* = 6, 12Hz, 2H, $CH_2$), 3.22 (t, *J* = 6Hz, 4H, 2 x $CH_2$), 3.43 (m, 1H, CH), 3.45-3.51 (m, 4H, piperidyl 2 x CH2), 3.62 (m, 2H), 3.63-3.67 (m, 4H, 2 x $CH_2$), 7.31 (d, *J* = 6 Hz, 1H, ArH), 8.03 (d, *J* = 12 Hz, 1H, ArH), 8.81 (s, 1H, ArH), 10.1 (brs, 1H, NH). $^{13}$C-NMR: ($CDCl_3$) δ 8.2, 25.4, 26.8, 28.4, 29.5, 32.6, 34.7, 39.2, 45.1, 50.0, 70.0, 70.2, 70.6, 71.3, 105.0, 111.4, 112.7, 112.8,122.2, 138.4, 144.8, 146.8, 152.6, 154.2, 154.6, 165.1, 175.4.

Synthesis of compound **(7)**:

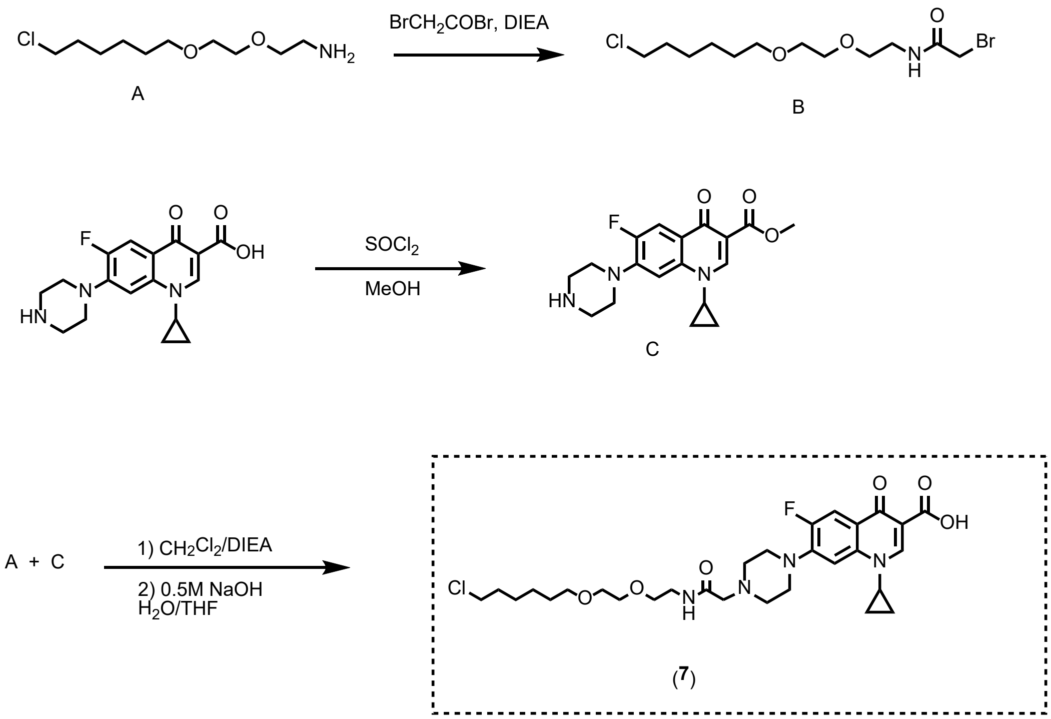


Compound (**7**). **B**: A solution of HaloTag amine ligand (26.0mg, 0.1mmol) was dissolved in 2.0mL of dry methylene chloride. The solution was cooled on an ice water bath, and to this solution was added diisopropylethylamine(61μL, 46,5mg, 0.36mmol). In another vial, bromoacetyl bromide (40.0mg, 0.2mmol) was dissolved in 1.0mL methylene chloride and cooled in an ice bath, and to the bromoacetyl bromide solution was added (50μL, 0.36mmol) of triethylamine via a syringe. This mixture was stirred for 30 minutes, and it was then added to an ice-cold water solution of the HaloTag amine ligand prepared as described above. The reaction mixture was stirred on an ice water bath for 30 minutes, allowed to warm to room temperature, and was further stirred for 1hr and concentrated under reduced pressure. The crude residue was diluted with 15 mL of ether, and to this, 10% $NaHCO_3$ (5.0mL) was added, stirred for 5 min and transferred to a separatory funnel to remove the aqueous layer. The organic ether layer was washed with dilute HCl, followed by sat. $NaHCO_3$. The ether layer was finally washed with brine, dried over $Na_2SO_4$, concentrated, and used as is for the next reaction (31.0mg).

Synthesis of ciprofloxacin methyl ester, **C** was carried out according to a reported procedure.[1] The compound was characterized by mass spectroscopy and $^1$H-NMR spectroscopy. MF: $C_{19}H_{22}FN_3O_3$, Calc. 346.15; Obs. 346.1 for $[M+H]^+$. $^1$H-NMR: ($CDCl_3$) $\delta$ 1.14 (m, 2H, cyclopropyl-$CH_2$), 1.32 (m, 2H, cyclopropyl-$CH_2$), 3.12 (m, 4H, piperidyl 2 x $CH_2$), 3.27 (m, 4H, piperidyl 2 x $CH_2$), 3.43 (m, 1H, cyclopropyl CH-N), 3.91 (s, 3H, $OCH_3$), 7.26 (d, *J* = 6 Hz, 1H, ArH), 8.02 (d, *J* = 18 Hz, 1H, ArH), 8.54 (s, 1H, ArH).

The reaction of bromoacetyl-HaloTag amide with ciprofloxacin methyl easter, **A + C**: The ciprofloxacin methyl ester (18.0mg, 0.052mmol) was dissolved in methylene chloride and to this, bromeacetyl-HaloTag amide (25.0mg, 0.072mmol, 1.4 eq) was added followed by 40 $\mu$L of DIEA (0.23 mmol). The mixture was stirred overnight at room temperature and the reaction was analyzed by TLC. A new spot having fluorescence was observed (more non- polar compared with the starting ciprofloxacin methyl ester). The compound was purified by silica gel column chromatography using hexane/ethyl acetate (1:1) as eluent. The homogenous fractions as observed by TLC were combined and concentrated to afford ciprofloxacin-N-alkylated HaloTag methyl ester (23.0mg, yield: 73%). $^1$H-NMR: ($CDCl_3$) $\delta$ 0.92 (dd, *J* = Hz, 2H, Cyclopropyl CH2), 1.20-1.39 (m, 18H), HRMS: for MF $C_{30}H_{43}ClFN_4O_6$, Calc. 609.2850; Obs. 609.2846 for $[M+H]^+$.

Hydrolysis of the methyl ester to afford Compound **(7)**: The ciprofloxacin-N-alkylated HaloTag amide (20.0mg, 0.033mmol) was dissolved in THF (5.0mL) and to this was added 1.0M NaOH (200 μL). The mixture was vigorously stirred at room temperature for 4 hrs. TLC analysis showed that the methyl ester was consumed to form a new non-polar compound. The reaction mixture was concentrated under reduced pressure, acidified with dil. HCl (1M) and extracted with methylene chloride. The organic layer was washed, dried over $Na_2SO_4$, and evaporated under reduced pressure to yield the crude residue which was purified with silica column chromatography using ($CHCl_3$/MeOH; 98:2) as eluent. Homogenous fractions as observed by TLC were combined and concentrated under reduced pressure to yield a white solid (14.5 mg, yield: 74%). $^1$H-NMR: ($CDCl_3$) δ 1.14 (m, 2H, cyclopropyl-$CH_2$), 1.32 (m, 2H, cyclopropyl-$CH_2$), 1.39-1.49 (m, 6H, 3 x $CH_2$), 1.49 (s, 9H, 3 x$CH_3$), 1.58 (dt, *J* = 6Hz, 128Hz, 2H, $CH_2$), 1.74 (dt, *J* = 6, 12Hz, 2H, $CH_2$), 3.22 (t, *J* = 6Hz, 4H, 2 x $CH_2$), 3.43 (m, 1H, CH), 3.45-3.51 (m, 4H, piperidyl 2 x CH2), 3.62 (m, 2H), 3.63-3.67 (m, 4H, 2 x $CH_2$), 7.31 (d, *J* = 6 Hz, 1H, ArH), 8.03 (d, *J* = 12 Hz, 1H, ArH), 8.81 (s, 1H, ArH), 10.1 (brs, 1H, NH). $^{13}$C-NMR: ($CDCl_3$) δ 8.2, 25.4, 26.8, 28.4, 29.5, 32.6, 34.7, 39.2, 45.1, 50.0, 70.0, 70.2, 70.6, 71.3, 105.0, 111.4, 112.7, 112.8, 122.2, 138.4, 144.8, 146.8, 152.6, 154.6, 165.1, 175.4. HRMS: for MF; $C_{29}H_{40}ClFN_4O_6$ Calc.. 595.2699; Obs. 596.2690 for $[M+H]^+$.

Synthesis of compound **(8)**:

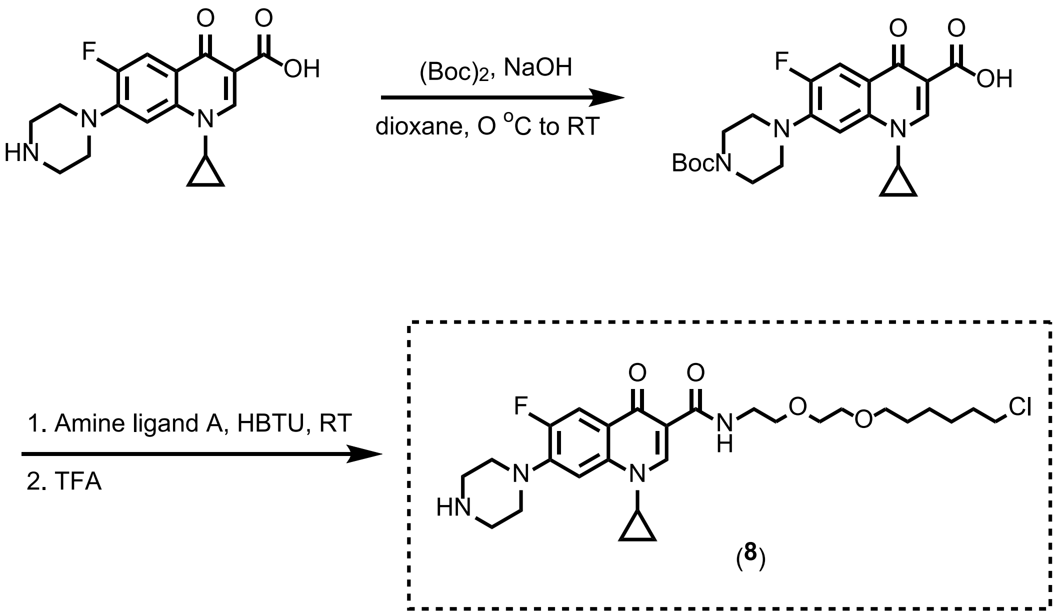


Compound (**8**). **N-$^t$Boc-Ciprofloxacin Synthesis:** The compound was synthesized using a reported procedure with slight modifications.[2] Briefly, ciprofloxacin (330.0mg, 1 mmol) was suspended in 25 mL of 1,4-dioxane. 10 mL of 1.0 M NaOH was added to the suspension and the mixture was stirred on an ice bath for 30 minutes prior to addition of di-tert-butyl dicarbonate (250mg, 1.15 mmol). The mixture was allowed to warm up to room temperature and stirred overnight. Concentration of volatiles under reduced pressure using a rotary evaporator afforded a white solid which was filtered, washed with ice-cold water and with ether. The solid was dried in air to yield 7-(4-(tert-butoxycarbonyl)piperazin-1-yl)-1-cyclopropyl-6-fluoro-4-oxo-1,4-dihydroquionoline-3-carboxylic acid (375.0 mg. 87%). $^1$H-NMR and mass spectral data matched with the reported compound.

**Reaction of N-$^t$Boc-ciprofloxacin with HaloTag amine ligand, synthesis of N-$^t$Boc-ciprofloxacin-carboxy HaloTag amide:** N-$^t$Boc-ciprofloxacin (86.2 mg, 2mmol) was dissolved in 2.0 mL anhydrous DMF and to this was added DIEA (50μL) followed by HBTU (100.0mg, 2.6 mmol). After stirring for 1hr at room temperature, a mixture of HaloTag amine (60.0mg, 2.3 mmol) and DIEA (100 μL) in 0.5 mL DMF was added to the solution and further stirred for 4 hrs. TLC analysis indicated complete loss of the starting material to form a non-polar compound. Dichloromethane (25 ml) was added to the reaction mixture along with 15.0 mL of water transferred to a separatory funnel. The dichloromethane layer was isolated and washed with dil. HCl, water, and finally with brine. The organic layer was then dried over $Na_2SO_4$ and concentrated under reduced pressure yielding a residue which was further purified with flash chromatography over silica gel ($CHCl_3$/MeOH, 98:2). The homogenous fractions were collected and concentrated to yield a white solid (98.0mg, yield: 77 %). $^1$H-NMR: ($CDCl_3$) δ 1.14 (m, 2H, cyclopropyl-$CH_2$), 1.32 (m, 2H, cyclopropyl-$CH_2$), 1.39-1.49 (m, 6H, 3 x $CH_2$), 1.49 (s, 9H, 3 x$CH_3$), 1.58 (dt, *J* = 6Hz, 128Hz, 2H, $CH_2$), 1.74 (dt, *J* = 6, 12Hz, 2H, $CH_2$), 3.22 (t, *J* = 6Hz, 4H, 2 x $CH_2$), 3.43 (m, 1H, CH), 3.45-3.51 (m, 4H, piperidyl 2 x CH2), 3.62 (m, 2H), 3.63-3.67 (m, 4H, 2 x $CH_2$), 7.31 (d, *J* = 6 Hz, 1H, ArH), 8.03 (d, *J* = 12 Hz, 1H, ArH), 8.81 (s, 1H, ArH), 10.1 (brs, 1H, NH). $^{13}$C-NMR: ($CDCl_3$) δ 8.2, 25.4, 26.8, 28.4, 29.5, 32.6, 34.7, 39.2, 45.1, 50.0, 70.0, 70.2, 70.6, 71.3, 105.0, 111.4, 112.7, 112.8, 122.2, 138.4, 144.8, 146.8, 152.6, 154.6, 165.1, 175.4.

**Deprotection of N-tBoc-ciprofloxacin-carboxy HaloTag amide:** N-$^t$Boc-ciprofloxacin-carboxy HaloTag amide (60.0mg, 0.094 mmol) was dissolved in 3.0mL of dichloromethane in a 20 mL RB flask and stirred on an ice bath, and to this was added 0.5 mL of TFA. The mixture was stirred for 1hr, during which time TLC analysis indicated complete consumption of the starting material. The reaction mixture was concentrated under reduced pressure and dichloromethane (5.0mL) was added and evaporated. This dichloromethane cycle was repeated one more time to remove TFA to afford Ciprofloxacin-carboxy-HaloTag amide TFA salt as Compound **(8)**. Trituration with ether gave a white solid (57.0 mg). $^1$H-NMR: ($CDCl_3$) $\delta$ 1.17 (m, 2H, cyclopropyl-$CH_2$), 1.34-1.42 (m, 4H, cyclopropyl-$CH_2$ & $CH_2$), 1.43 (m, 2H, $CH_2$), 1.58 (dt, $J$ = 6Hz, 128Hz, 2H, $CH_2$), 1.73 (dt, $J$ = 6, 12Hz, 2H, $CH_2$), 3.16 (m, $CH_2$), 3.45-3.55 (m, 12H, $CH_2$), 3.55 (m, 2H, $CH_2$), 3.60-3.69 (m, $CH_2$), 7.37 (d, $J$ = 6 Hz, 1H, ArH), 8.04 (d, $J$ = 12 Hz, 1H, ArH), 8.82 (s, 1H, ArH), 10.1 (brs, 1H, NH). $^{13}$C-NMR: ($CDCl_3$) $\delta$ 8.3, 25.5, 26.8, 29.6, 32.7, 34.9, 39.3, 43.6, 45.2, 47.1, 50.0, 70.0, 70.2, 70.6, 71.4, 111.6, 113.1, 123.3, 138.4, 143.4, 143.5, 147.2, 152.5, 154.1, 175.3.

**Biochemical Methods.** For all biological assays, laboratory strains of wild type *E. coli* (ATCC 25922), and K12 *E. coli* strains BL21(DE3) and *E. coli* Lemo21(DE3) were grown with shaking at 37 °C overnight from freezer glycerol stocks in 5 mL of Luria Broth (LB) media. mc$^2$155 *M. smegmatis* (NC_008596 in GenBank) and H37Rv Δ*panCD* Δ*leuCD* *Mycobacterium tuberculosis* strain mc$^2$6206 were grown at 37°C in shaking in Middlebrook 7H9 growth medium (BD Difco, Franklin Lakes, NJ) supplemented with 0.4% glycerol, 0.05% Tween-80 and 10% ADC for *M. smegmatis* or OADC, 50 μg/ml pantothenic acid and 50 μg/ml L-leucine for *M. tuberculosis*.

**HaloTag Protein Expression: Cytoplasmic HaloTag protein.** For expression of the HaloTag protein in *E. coli*, HaloTag expression plasmid Cyto_Halo_pET21(b)+ was transformed into *E. coli* BL21(DE3) and grown on an LB/agar plate with ampicillin (100 μg/mL) at 37ºC overnight. Colonies were picked and grown in 5 mL LB broth with ampicillin at 37ºC overnight. A culture tube containing 2 mL LB media was inoculated with the overnight culture at a ratio of 1:100 and cells were grown at 37 ºC for 3 hours, or until the optical density at 600 nm reached 0.6. Cultures were induced with 1 mM Isopropyl β-d-1-thiogalactopyranoside (IPTG) at 37°C for 2 hours in a shaker incubator, and then pelleted at 4,000 rpm for 3 min in an HERAEUS Multicentrifuge X1 centrifuge (Thermofisher Scientific). The pellet was washed three times with 1X PBS pH 7.2 following which cells were ready for assaying. The identity of the protein was confirmed by SDS-PAGE.

**HaloTag Protein Expression: Periplasmic HaloTag protein.** For localization of HaloTag protein in the periplasm, a modification of Cyto_Halo containing an amine terminal recognition sequence, DsbA for export to the periplasm by the SecA pathway, i.e., Peri_Halo_pET21(b)+ was transformed into *E. coli* Lemo21(DE3) and grown on an LB/agar plate with ampicillin (100 μg/mL) and chloramphenicol (34μg/mL) at 37 ºC overnight. Colonies were picked and grown in 5 mL LB broth with ampicillin/chloramphenicol at 37 ºC overnight. A culture tube containing 2mL LB media was inoculated with the overnight culture at a ratio of 1:100 and cells were grown at 37 ºC for 3hrs, or until the optical density at 600 nm reached 0.6. Cultures were induced with 1 mM Isopropyl β-d-1-thiogalactopyranoside (IPTG) at 37°C for 2 hours in a shaker incubator, and then pelleted at 4,000 rpm for 3 min in an HERAEUS Multicentrifuge X1 centrifuge (Thermofisher Scientific). The pellet was washed three times with 1X PBS pH 7.2 following which cells were ready for assaying. The identity of the protein was confirmed by SDS-PAGE.

**BaCAPA Pulse-Chase Protocol (chloroalkane-tagged molecule, chloroalkane-tagged dye).** For a bacterial chloroalkane penetration assay (BaCAPA), 25 μL of serially diluted samples of the chloroalkane-tagged test molecule, in quadruplicate, were transferred into a 96-well plate at 2X of the desired concentration. The pellet obtained from the washing step above was resuspended in 2mL of 1X PBS by vortexing thoroughly. 25 μL of the cell suspension (both IPTG induced and non-induced cells was added into the wells and incubated at 37°C with shaking at 250 RPM. After 30 min, the plate was centrifuged to pellet the cells using a Jouan C4i centrifuge (Thermofisher Scientific). The supernatant was discarded, and the pellet was washed twice with PBS. The chase step followed by addition into each test well, 50 μL of 5 μM chloroalkane-tagged rhodamine 110 and incubated for 30 min at 37°C with shaking in a shaker incubator at 250 RPM. The plate was centrifuged to pellet the cells and cells were washed three times with PBS and fixed with 4% formaldehyde by incubating at room temperature for 30 min. The cells were analyzed by flow cytometry on the Attune NxT Acoustic Focusing Cytometer (Invitrogen) by exciting using the blue laser (530nm).

**BaCAPA Control Experiment: pulsing with a chloroalkane-tagged dye.** 25 μL of the cell suspension (both IPTG induced and non-induced cells) was added into the wells of a 96-well plate. The plate was centrifuged to pellet the cells using a Jouan C4i centrifuge (Thermofisher Scientific). The supernatant was discarded and into each test well, 50 μL of chloroalkane-tagged rhodamine 110 was added and incubated for 30 min at 37°C in a shaker incubator (250 RPM). The plate was centrifuged to pellet the cells and cells were washed three times with PBS and fixed with 4% formaldehyde by incubating at room temperature for 30min. The cells were analyzed by flow cytometry on the Attune NxT Acoustic Focusing Cytometer (Invitrogen) by exciting using the blue laser (530 nm).

**Polymyxin nonapeptide (PMBN) mediated bacterial chloroalkane-tagged molecule penetration.** For the PMBN-mediated penetration assay, 25 μL of the desired PMBN concentration was added into the wells of a 96-well plate at 3X the final concentration along with 25 μL of the HaloTag protein cell suspension (both IPTG induced and non-induced cells). Next, 25 μL of the serially diluted samples of the chloroalkane-tagged test molecule was added into the wells at 3X the desired final concentration in quadruplicate. The plates were incubated at 37°C with shaking at 250 RPM. After 3 0min, the plate was centrifuged to pellet the cells using a Jouan C4i centrifuge (Thermofisher Scientific). The supernatant was discarded, and the pellet was washed twice with PBS. The chase step followed by addition into each test well, 50 μL of 5 μM chloroalkane-tagged rhodamine 110 and incubated for 30 min at 37°C with shaking in a shaker incubator at 250 RPM. The plate was centrifuged to pellet the cells and cells were washed three times with PBS and fixed with 4% formaldehyde by incubating at room temperature for 30 min. The cells were analyzed by flow cytometry on the Attune NxT Acoustic Focusing Cytometer (Invitrogen) by exciting using the blue laser (530 nm).

**Mammalian Cell Culture.** J774A.1 cells were cultured in Dulbecco's modified Eagle's medium (DMEM) supplemented with 10% (v/v) FBS, 50 IU/mL penicillin, 50 μg/mL streptomycin, and 2 mM L-glutamine in a humidified atmosphere of 5% $CO_2$ at 37°C.

**BaCAPA labeling of *E. coli* inside macrophages.** J774A.1 macrophages were cultured to confluency, resuspended in antibiotic free media (DMEM + 10% FBS containing no Pen-Strep), and were added to a 96-well plate with 200,000 cells/well. The cells were centrifuged at 1,000 rpm for 5 mins to pellet the cells. The macrophage pellets were resuspended in antibiotic free media containing HaloTag protein expressing *E. coli* BL21(DE3) (either IPTG induced or non-induced) at a MOI 50 and incubated at 37°C for 15 mins to promote bacterial cell uptake into macrophages. The macrophages were pelleted, resuspended in DMEM + 10% FBS + 300 μg/mL gentamycin (to clear nonphagocytosed bacteria) + 5 μM of chloroalkane-tagged rhodamine 110 and incubated for 5 min at 37°C. The macrophages were washed 3x with 1X PBS to remove excess dye and cells were fixed for 10 min at 4°C with 4% formaldehyde in 1X PBS. Cells were analyzed using the Attune NxT Flow Cytometer (Thermo Fischer) equipped with a 488 nm laser and 525/40 nm bandpass filter. Cells were gated to analyze only the population of macrophages. For confocal microscopy, J774A macrophages were treated with 5 μg/mL of TMR-tagged Wheat Germ Agglutinin (Vector Laboratories, RL-1022) for 30 min at 4°C. Glass microscope slides were spotted with a 1% agar pad and 5 μL of sample were deposited onto the agar. Samples were covered with a micro cover glass and imaged using a Zeiss 880/990 multiphoton Airyscan microscopy system (40x oil-immersion lens) equipped with 488 nm and 550 nm lasers. Images were obtained and analyzed via Zeiss Zen software. We acknowledge the Keck Center for Cellular Imaging and for the usage of the Zeiss 880/980 multiphoton Airyscan microscopy system (PI- AP: NIH-OD025156).

**BaCAPA labeling of *E. coli* for confocal microscopy imaging.** 25 μL of the cell suspension of HaloTag protein expressing *E. coli* BL21(DE3) (either IPTG induced or non-induced) was added into the wells of a 96-well plate. The plate was centrifuged to pellet the cells using a Jouan C4i centrifuge (Thermofisher Scientific). The supernatant was discarded and into each test well, 50 μL of chloroalkane-tagged rhodamine 110 was added and incubated for 30 min at 37°C in a shaker incubator (250 RPM). The plate was centrifuged to pellet the cells and cells were washed three times with PBS and fixed with 4% formaldehyde by incubating at room temperature for 30min. The cells were analyzed by flow cytometry on the Attune NxT Acoustic Focusing Cytometer (Invitrogen) by exciting using the blue laser (530 nm). For confocal microscopy, glass microscope slides were spotted with a 1% agar pad and 2 μL of fixed bacterial samples were deposited onto the agar. Samples were covered with a micro cover glass and imaged using a Zeiss 880/990 multiphoton Airyscan microscopy system (60x oil-immersion lens) equipped with a 488 nm laser. Images were obtained and analyzed via Zeiss Zen software. We acknowledge the Keck Center for Cellular Imaging and for the usage of the Zeiss 880/980 multiphoton Airyscan microscopy system (PI- AP: NIH-OD025156).

**Minimum Inhibitory Concentration.** Compounds were serially diluted 2-fold from stock solutions to yield 12 test concentrations in U-bottom, 96-well plates. Overnight cultures were used to inoculate LB media 1:100 and regrown to exponential phase, as determined by optical density recorded at 600 nm ($OD_{600}$). All cultures were diluted again to ca. $10^6$ CFU/mL in MH media, and 100 μL was inoculated into each well of a U-bottom 96-well plate (BD Biosciences, BD 351177) containing 100 µL of compound solution (in line with CLSI Standards).[4] Plates were incubated statically at 37 °C for 24 h upon which time wells were evaluated visually for bacterial growth. The MIC was determined as the lowest concentration of compound resulting in no bacterial growth visible to the naked eye, based on the majority of three independent experiments. Growth and sterility controls were conducted for each plate.

***HaloTag* gene cloning in pMV361 and pL5ptet0 vectors.** The *HaloTag* gene was synthesized as a *gBlock* by IDT (Integrated DNA Technologies) after codon optimization (with *IDT Codon Optimization Tool*) for protein expression in *M. tuberculosis*, containing the overlapping regions to be cloned in pMV361[1] or pL5ptet0[2] according to manufacturer instruction of Gibson Assembly Master Mix (New England Biolabs). The correct sequence of the plasmids was verified by whole plasmid sequencing (Plasmidsaurus).

***rodA* gene cloning in *pL5ptet0_HaloTag* vector.** *rodA* gene was amplified from pL5ptet0_*RodA-mRFP* plasmid[2] and cloned in pL5ptet0_*HaloTag* plasmid frame with HaloTag to obtain pL5ptet0_*RodA-HaloTag*, using Gibson Assembly Master Mix (New England Biolabs). To ensure the functionality of the two proteins, *rodA* and *HaloTag* genes were cloned separated. The correct sequence of the plasmid was verified by whole plasmid sequencing (Plasmidsaurus).

**HaloTag labelling in *M. smegmatis.*** Early log-phase *M. smegmatis* ($OD_{600}$ 0.2-0.3) was treated by direct addition to the culture of chloroalkane fluorophores with the following concentrations: **R110cl** 1.5 μM, **COMcl** 2.5 μM, for **TAMRAdirect** ligand 0.25 μM. Bacteria were incubated with the fluorophore for 15 minutes at 37°C in shaking (400 rpm), then washed two times with PBSTB (PBS supplemented with 0.05% Tween-20, 0.01% BSA), fixed for 10 minutes with 4% PFA and washed 2 times with PBSTB.

**HaloTag labelling in *M. tuberculosis.*** Early log-phase *M. tuberculosis* ($OD_{600}$ 0.2-0.3) was treated by direct addition to the culture of chloroalkane fluorophores with the following concentrations: **R110cl** 1.5 μM or **TAMRAdirect** ligand 0.25 μM. Bacteria were incubated with the fluorophore for 15 minutes at 37°C in shaking, then washed two times with PBSTrB (PBS supplemented with 0.05% Triton-X-100, 0.01% BSA), fixed for 4 hours with 4% PFA and washed 2 times with PBSTrB.

**Competition assay in *M. smegmatis.*** Early log-phase *M. smegmatis* ($OD_{600}$ 0.2-0.3) was treated by direct addition to the culture of +/- amine-chloroalkane competitor 100 μM, for 15 minutes at 37°C in shaking (400 rpm), then washed two times with PBSTB (PBS supplemented with 0.05% Tween-20, 0.01% BSA), and treated with **R110cl** 1.5 μM for 15 minutes at 37°C in shaking (400 rpm). After two washes in PBSTB bacteria were fixed for 10 minutes with 4% PFA and washed 2 times with PBSTB.

**Competition assay in *M. tuberculosis.*** Early log-phase *M. tuberculosis* ($OD_{600}$ 0.2-0.3) was treated by direct addition to the culture of +/- amine-chloroalkane (1) or ciprofloxacin-chloroalkane (7) competitors at various concentration (100 μM when used as single concentration, or 1, 5, 10, 50, and 100 μM for the construction of the permeability curve) for 15 minutes at 37°C in shaking. Bacteria were then washed two times with PBSTrB (PBS supplemented with 0.05% Triton-X-100, 0.01% BSA), and treated with **R110cl** (1.5 μM for single point competition or 1 μM for the permeability curve) for 15 minutes at 37°C in shaking. After two washes in PBSTrB bacteria were fixed for 4 hours with 4% PFA and washed 2 times with PBSTrB.

**Fluorescence Microscopy of *M. smegmatis* and *M. tuberculosis.*** Fixed bacteria were imaged either by conventional fluorescence microscopy (Nikon Eclipse E600,

Nikon Eclipse Ti or Zeiss Axioscope A1 with 100x oil objectives) and images were rendered using FIJI.[3]

**Flow cytometry analysis of *M. smegmatis* and *M. tuberculosis.*** After fixation and washing, bacteria were resuspended in filtered PBS and analyzed with BD DUAL LSRFortessa, UMass Amherst Flow Cytometry Core Facility. Data were analysed with FlowJo software.

**BaCAPA labeling of *M. tuberculosis* inside macrophages.** Immortalized bone marrow-derived macrophages (iBMDM, gift of Dr. Christopher Sassetti) were grown in Dulbecco's modified eagle's medium (DMEM, high glucose, GenClone) supplemented with 10% Fetal bovine serum (Gibco) 1% HEPES (Gibco), incubated at 37°C (atmosphere of 5% $CO_2$). The DMEM used during the infection was supplemented with 50 μg/ml pantothenic acid and 50 μg/ml L-leucine. The protocol was adapted from a prior work[4]. iBMDM were seeded on 8-chamber slides (30000 cells/well) and the day after were infected with multiplicity of infection 5:1 (*M. tuberculosis*:iBMDM). *M. tuberculosis* transformed with empty or HaloTag-containing pMV361 vector were grown for 48 hours in the presence of 100 μM HADA[5], then washed three times with PBSB (PBS supplemented with 0.1% BSA), and then resuspended in DMEM. The bacterial suspension was added to the 8-chamber slides wells and incubated for 4 hours. Then, bacteria were washed two times with PBSB and incubated with DMEM containing 200 μM amikacin for one hour to kill extracellular bacteria. Infected iBMDM were washed one time with PBSB and incubated overnight in DMEM. Cells were then washed twice with

PBSB and treated with **TMRcl** 1 μM in DMEM for 15 minutes, then washed three times with PBS prior to imaging.

NMR spectra

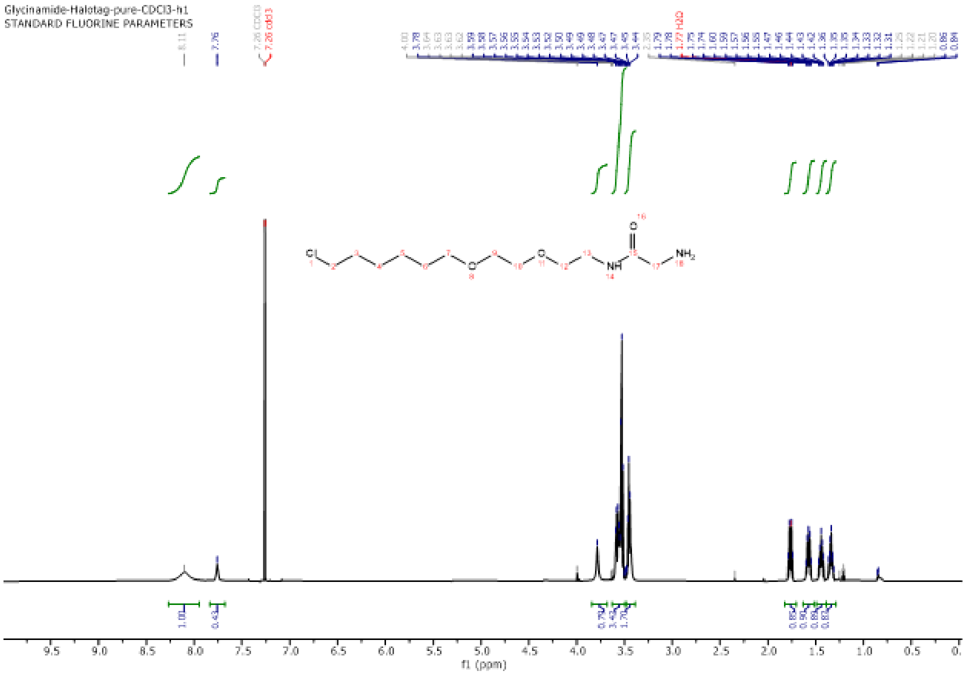
Glycinamide-Halotag-pure-CDCl3-h1
STANDARD FLUORINE PARAMETERS
f1 (ppm)

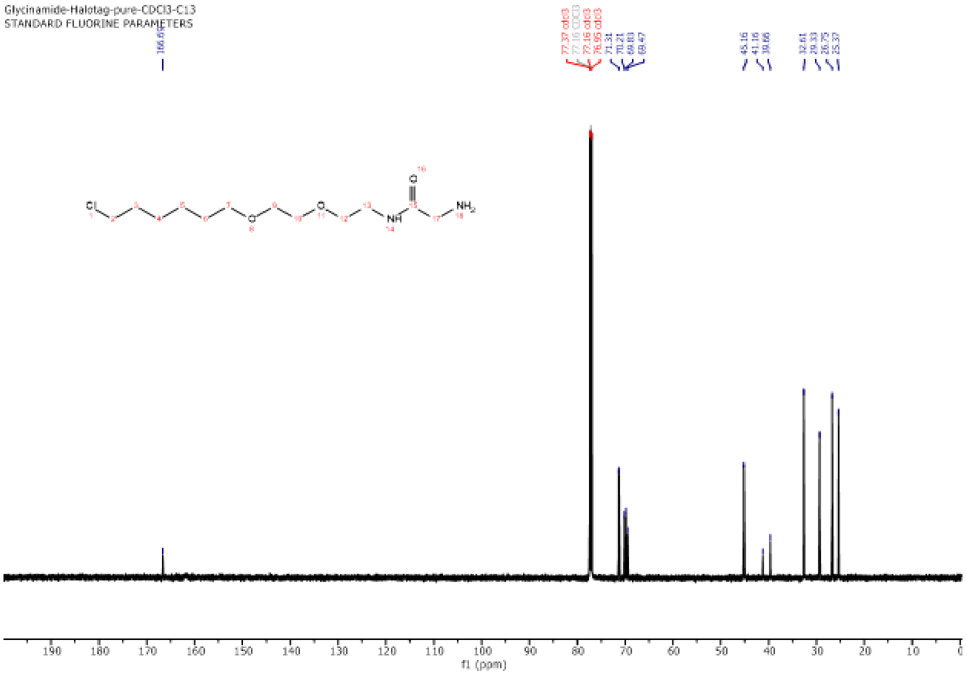
Glycinamide-Halotag-pure-CDCl3-C13
STANDARD FLUORINE PARAMETERS
166.69
77.37 cdcl3
77.16 CDCl3
77.16 cdcl3
76.95 cdcl3
71.31
70.21
69.83
69.47
45.16
41.16
39.66
32.61
29.53
26.75
25.37
Cl
NH
O
NH2
f1 (ppm)

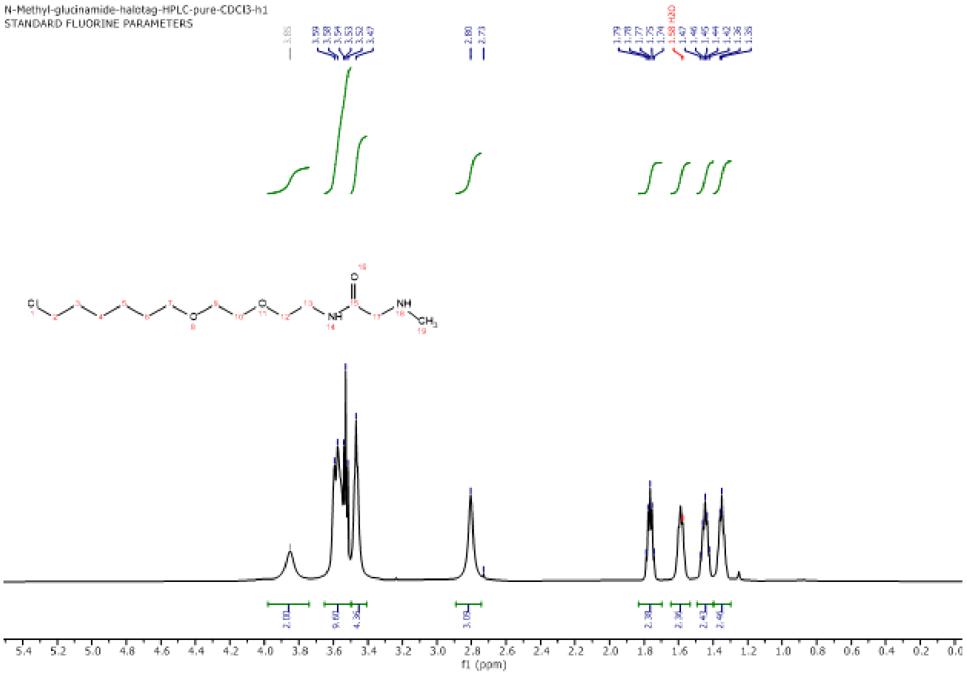
N-Methyl-glucinamide-halotag-HPLC-pure-CDCl3-h1
STANDARD FLUORINE PARAMETERS
f1 (ppm)

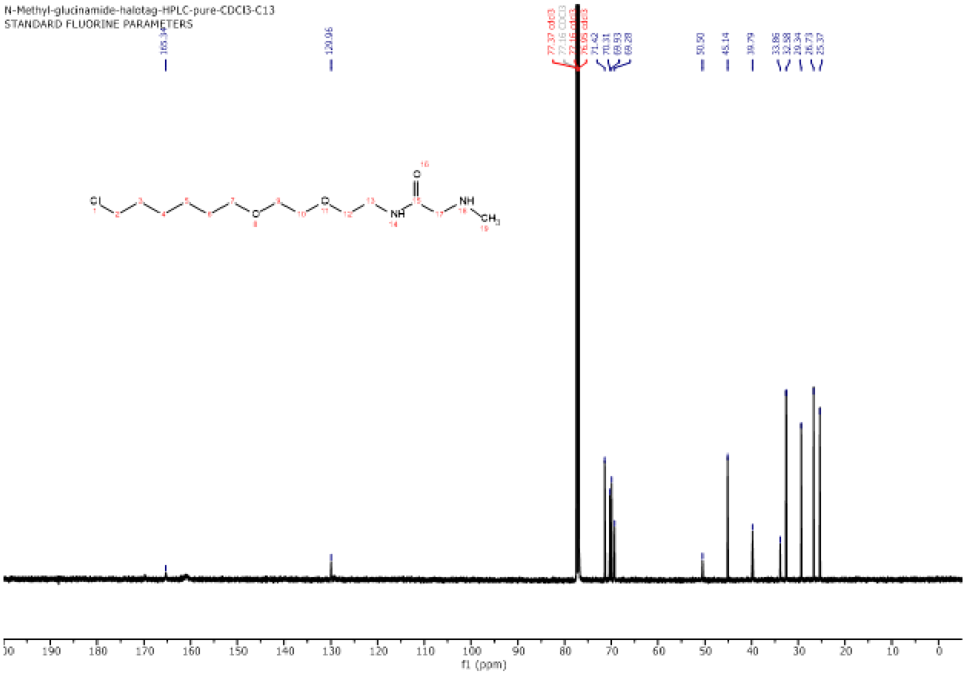

N-Methyl-glucinamide-halotag-HPLC-pure-CDCl3-C13
STANDARD FLUORINE PARAMETERS
165.34
129.96
77.37 cdcl3
77.16 CDCl3
77.16 cdcl3
76.95 cdcl3
71.62
70.31
69.93
69.28
50.50
45.14
39.79
33.86
32.88
29.34
26.73
25.37
f1 (ppm)

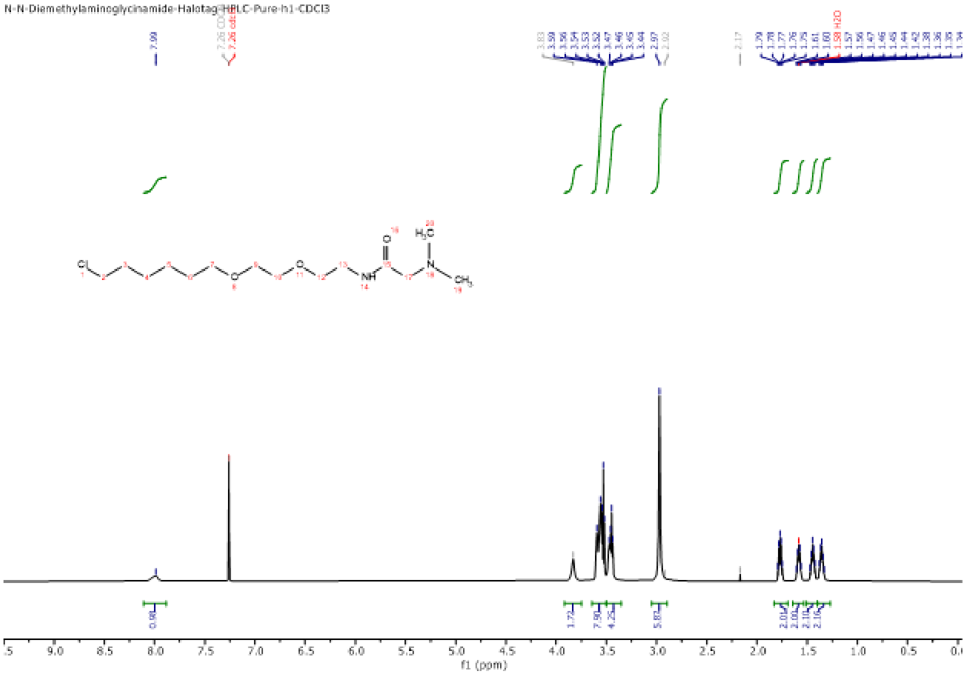
N-N-Diemethylaminoglycinamide-Halotag-HPLC-Pure-h1-CDCl3
f1 (ppm)

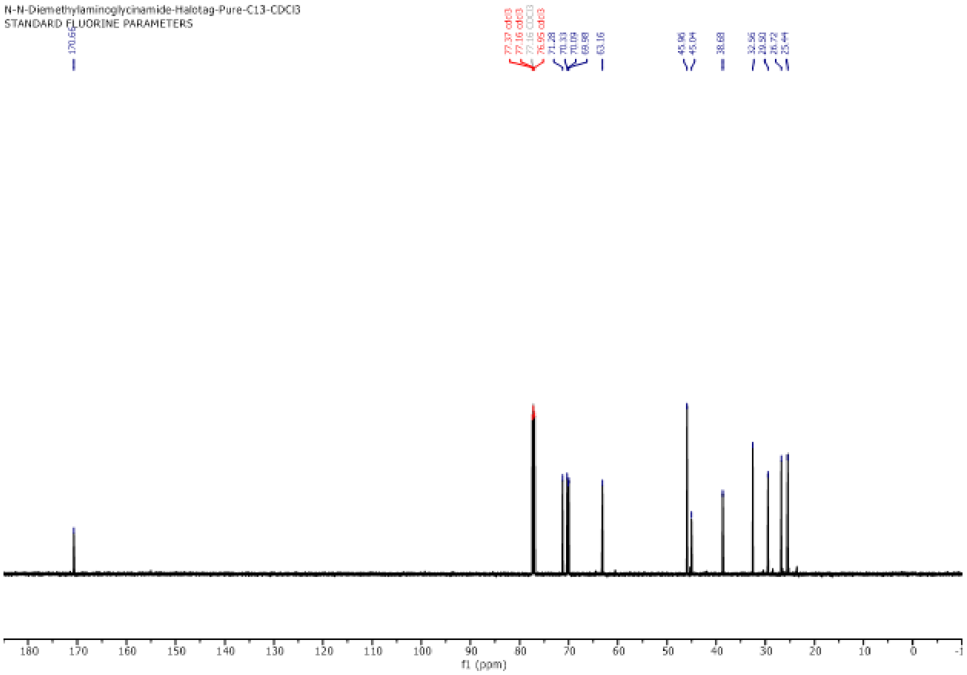
N-N-Diemethylaminoglycinamide-Halotag-Pure-C13-CDCl3
STANDARD FLUORINE PARAMETERS
170.66
77.37 cdcl3
77.16 cdcl3
77.16 CDCl3
76.95 cdcl3
71.28
70.33
70.09
69.98
63.16
45.96
45.04
38.68
32.56
29.50
26.72
25.44
180
170
160
150
140
130
120
110
100
90
80
70
60
50
40
30
20
10
0
f1 (ppm)

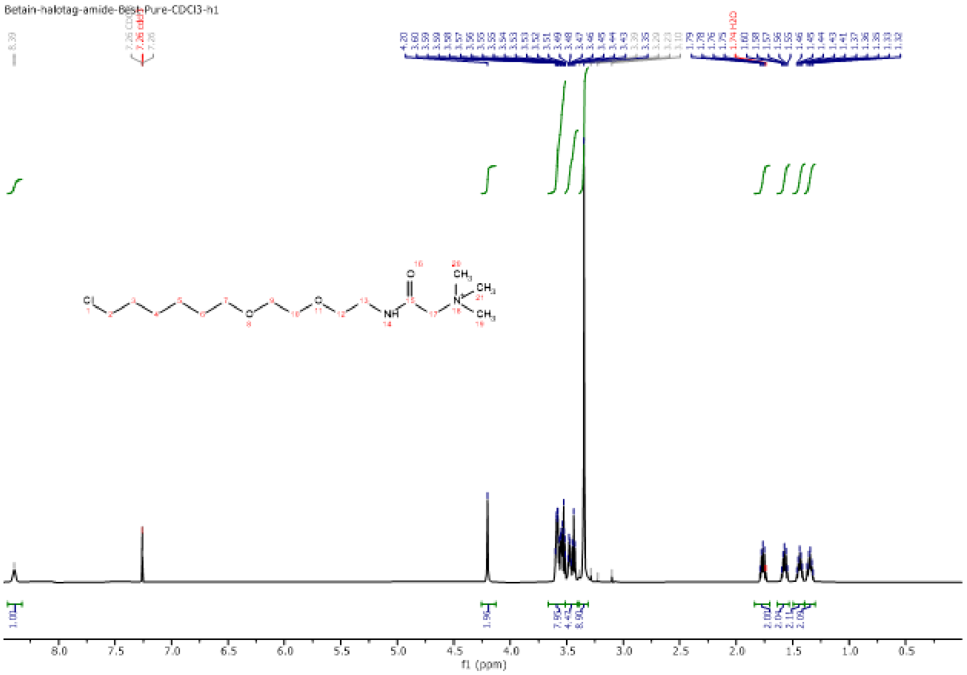

Betain-halotag-amide-Best-Pure-CDCl3-h1
f1 (ppm)

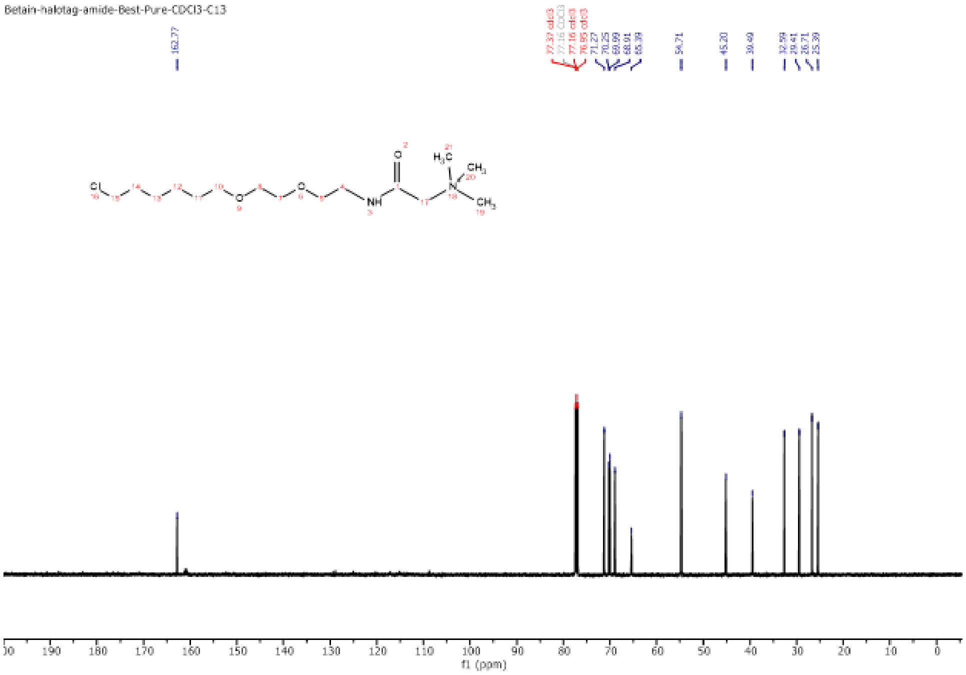
Betain-halotag-amide-Best-Pure-CDCl3-C13
162.77
77.37 cdcl3
77.16 CDCl3
77.16 cdcl3
76.95 cdcl3
71.27
70.25
69.99
68.91
65.39
54.71
45.20
39.49
32.59
29.41
26.71
25.39
f1 (ppm)

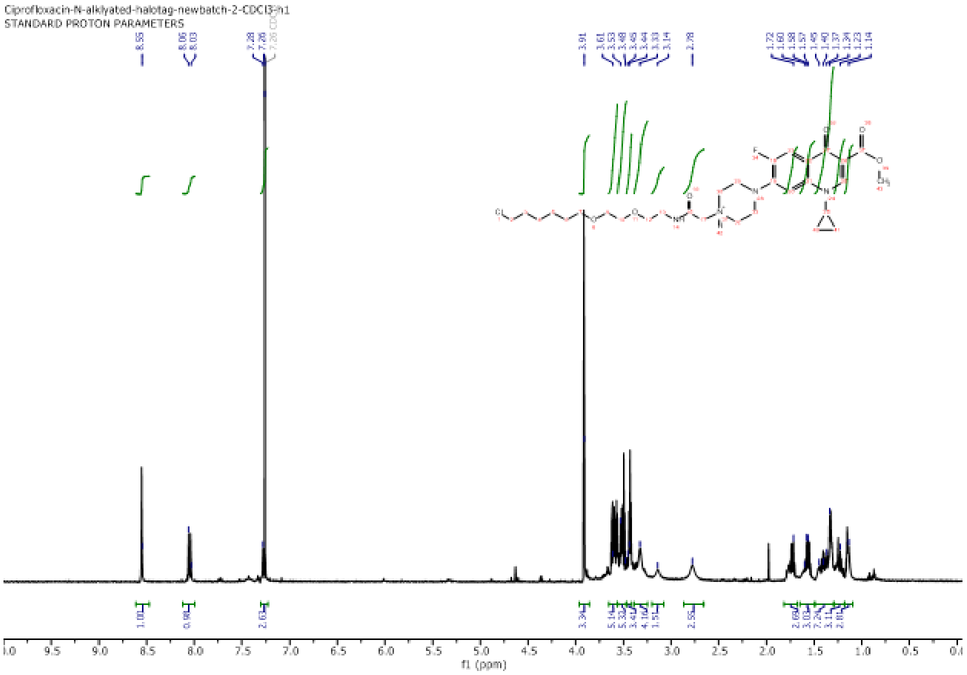
Ciprofloxacin-N-alklyated-halotag-newbatch-2-CDCl3-h1
STANDARD PROTON PARAMETERS
f1 (ppm)

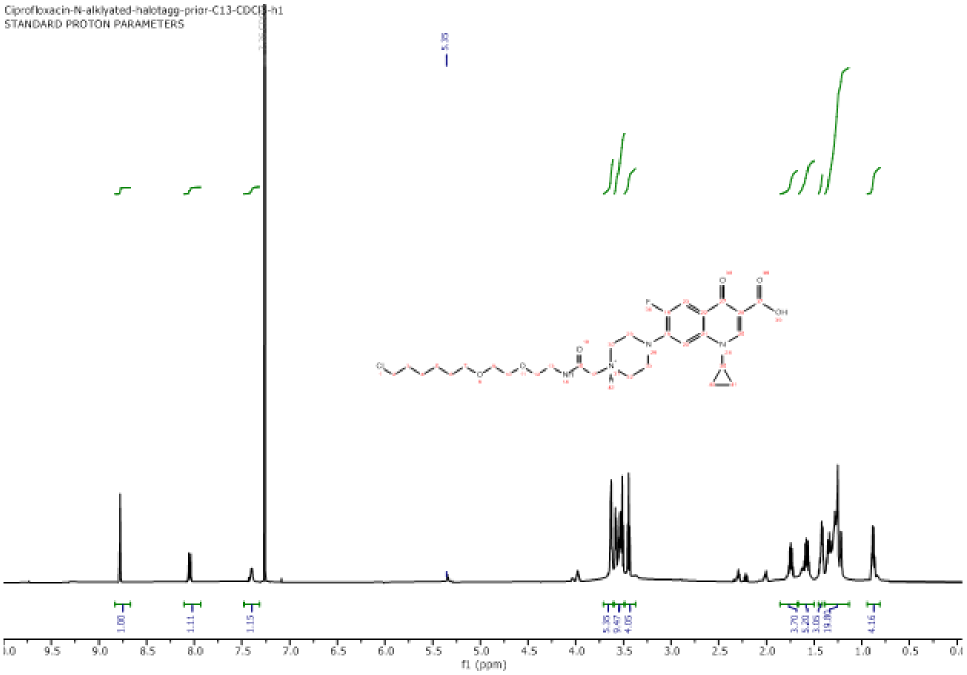
Ciprofloxacin-N-alklyated-halotagg-prior-C13-CDCl3-h1
STANDARD PROTON PARAMETERS
f1 (ppm)

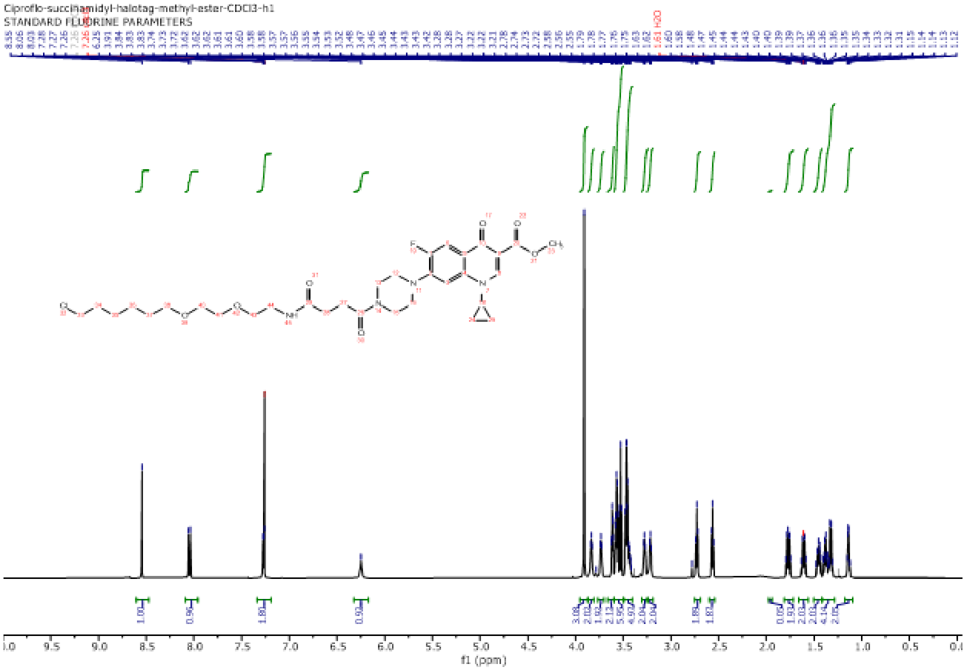

Ciproflo-succinamidyl-halotag-methyl-ester-CDCl3-h1
STANDARD FLUORINE PARAMETERS
f1 (ppm)

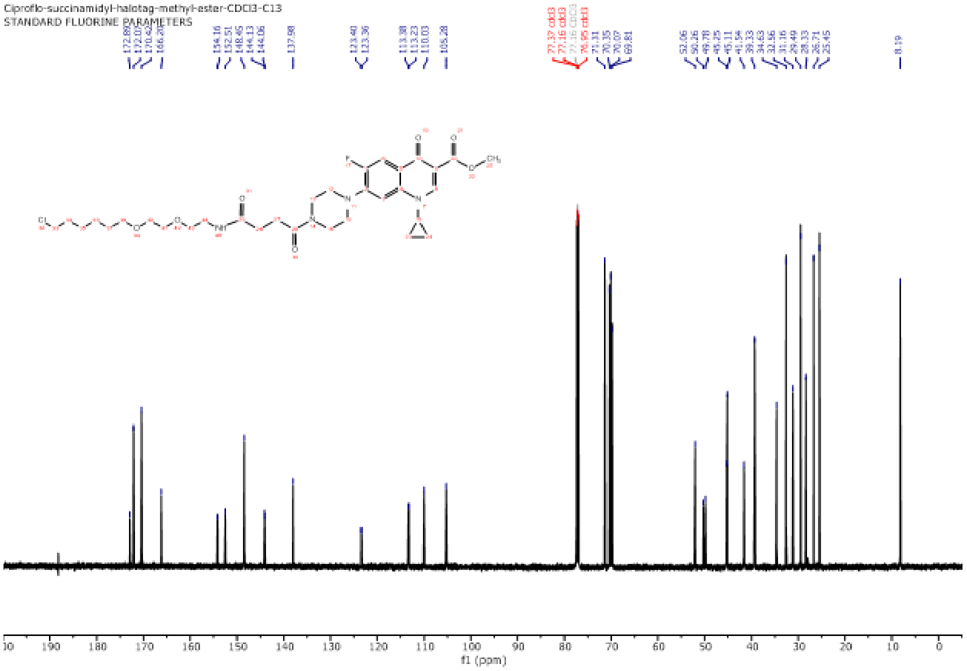
Ciproflo-succinamidyl-halotag-methyl-ester-CDCl3-C13
STANDARD FLUORINE PARAMETERS
f1 (ppm)

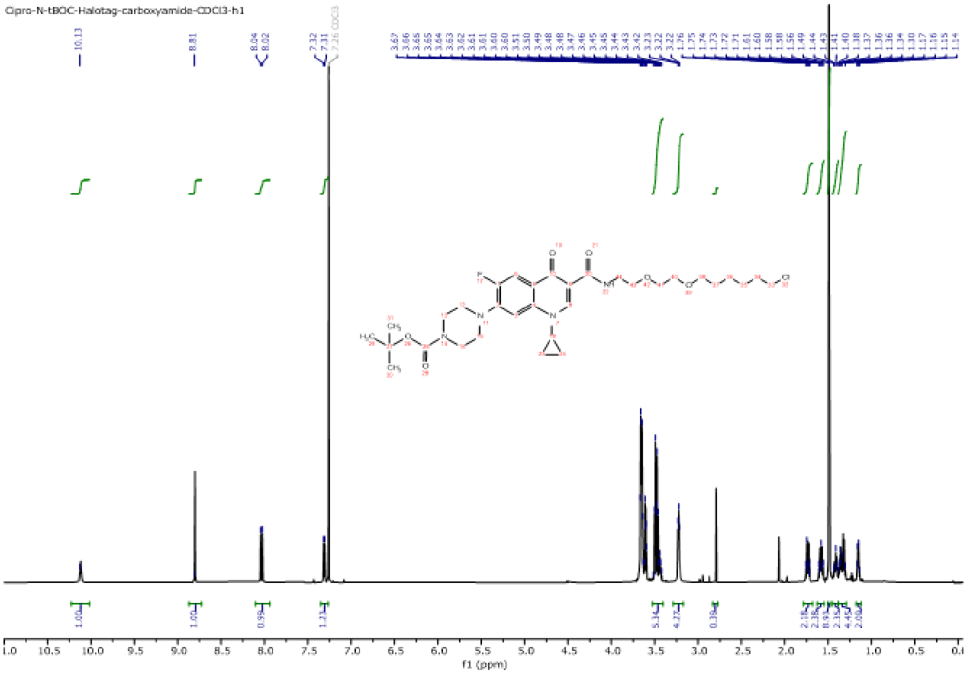
Cipro-N-tBOC-Halotag-carboxyamide-CDCl3-h1
f1 (ppm)

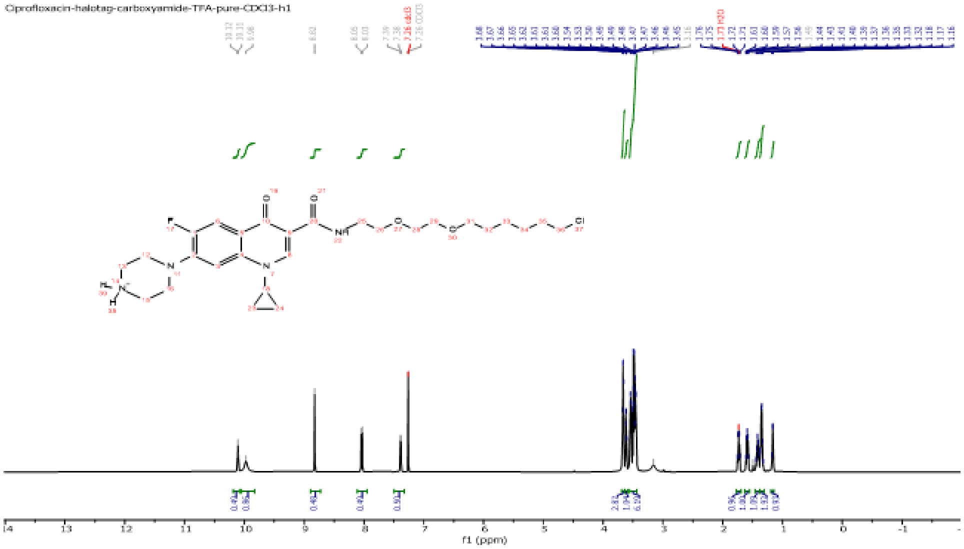

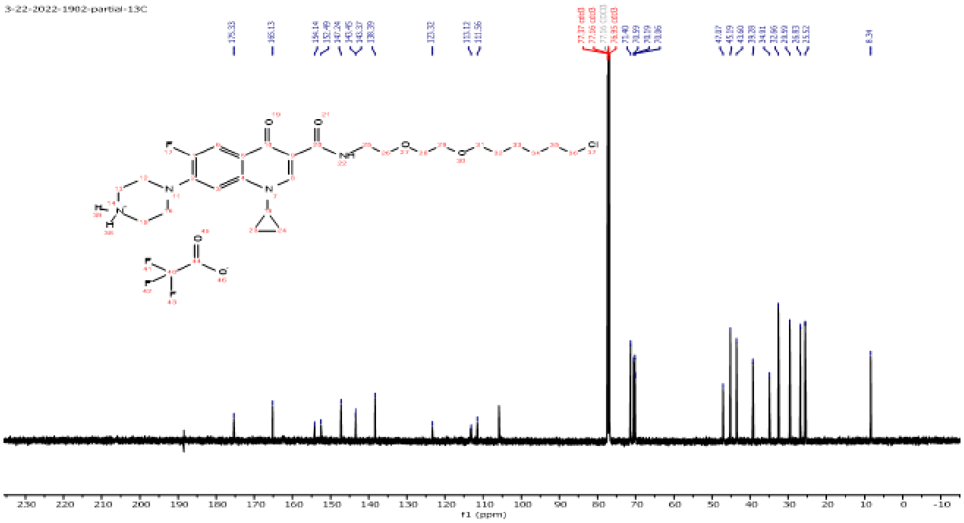
3-22-2022-1902-partial-13C
230 220 210 200 190 180 170 160 150 140 130 120 110 100 90 80 70 60 50 40 30 20 10 0 -10
f1 (ppm)

Analytical HPLC profiles:

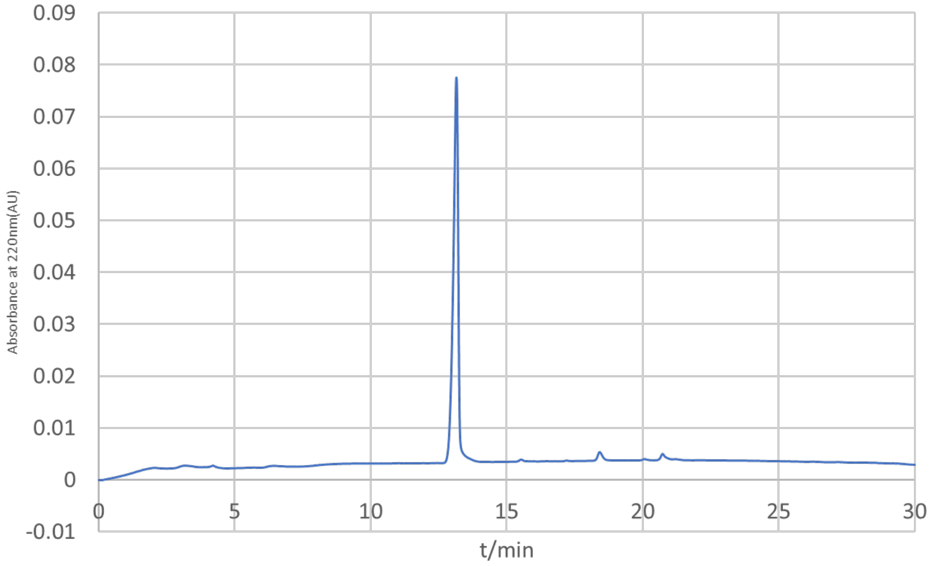


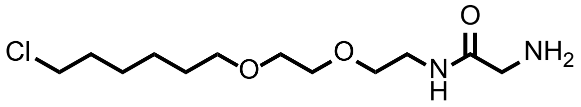


(**2**)

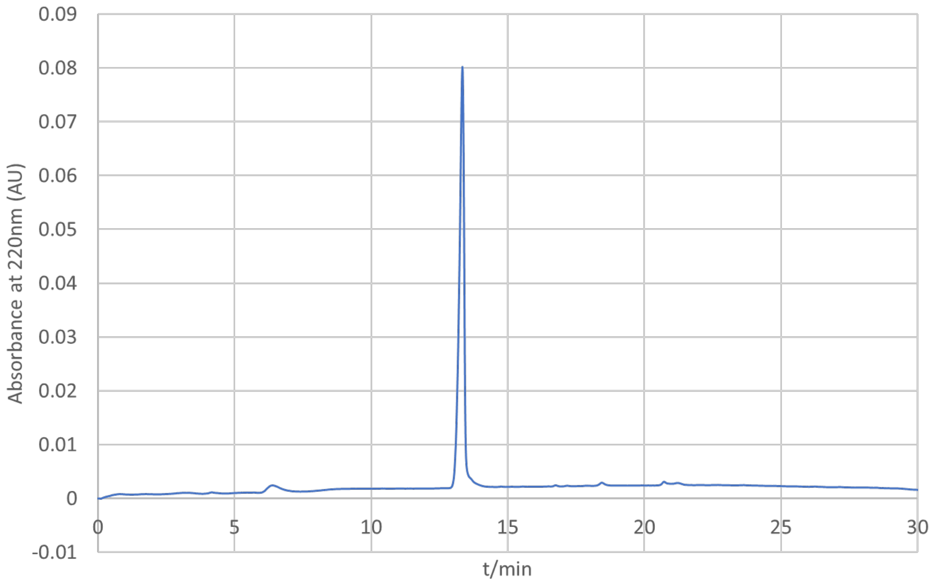

0.09
0.08
0.07
0.06
0.05
0.04
0.03
0.02
0.01
0
-0.01
Absorbance at 220nm (AU)
0
5
10
15
20
25
30
t/min


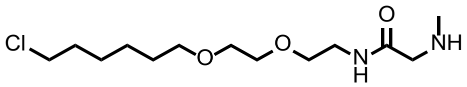

Cl
O
O
N
H
O
NH


(**3**)

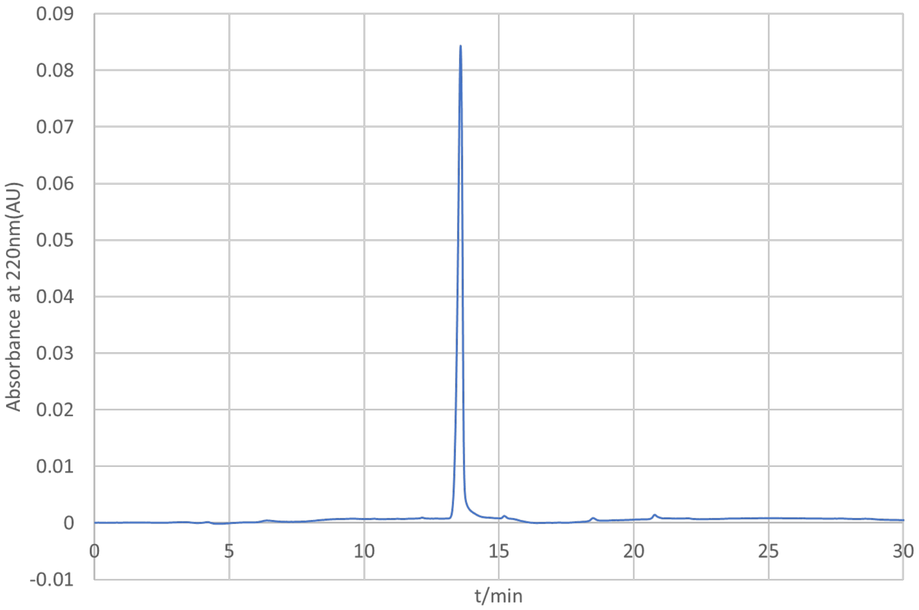

0.09
0.08
0.07
0.06
0.05
0.04
0.03
0.02
0.01
0
-0.01
Absorbance at 220nm(AU)
0
5
10
15
20
25
30
t/min


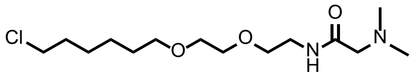

Cl
O
O
O
N
H
N


(**4**)

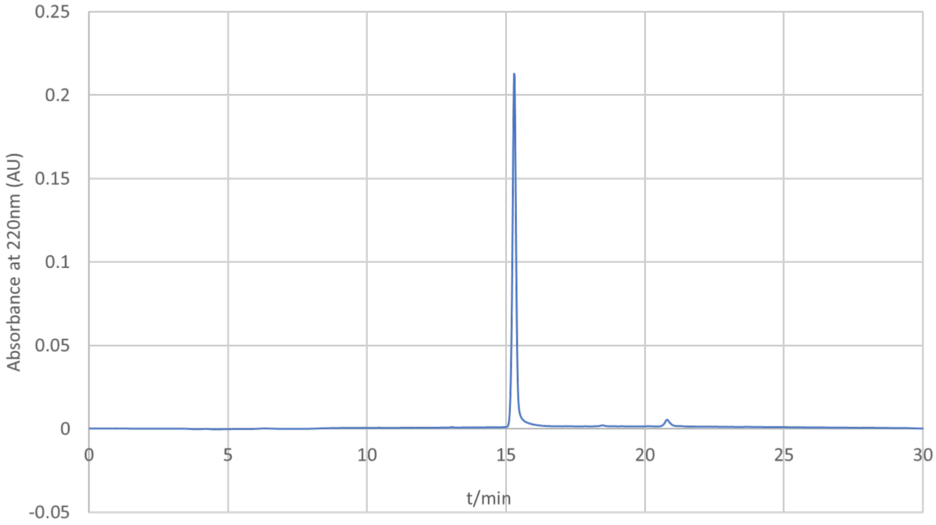
0.25
0.2
0.15
0.1
0.05
0
-0.05
Absorbance at 220nm (AU)
0
5
10
15
20
25
30
t/min

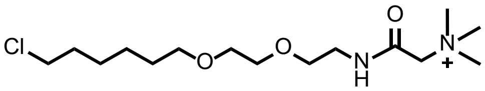
Cl
O
O
N
H
O
N
+

(**5**)

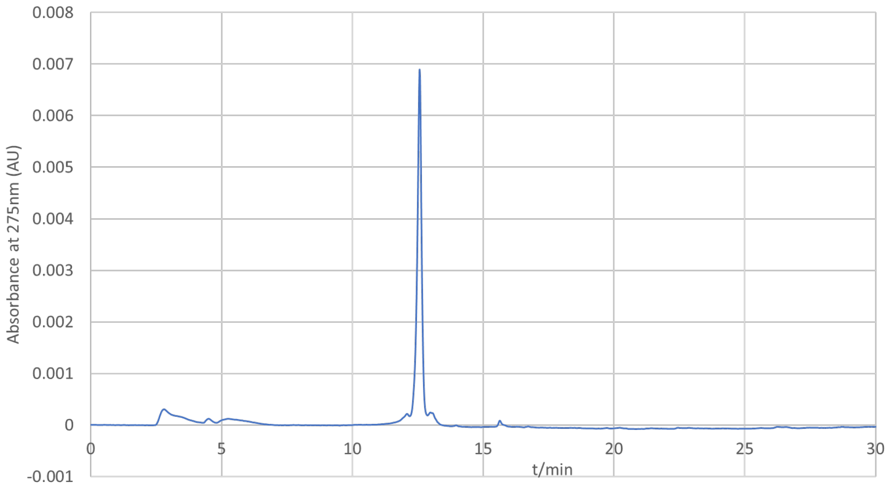
0.008
0.007
0.006
0.005
0.004
0.003
0.002
0.001
0
-0.001
Absorbance at 275nm (AU)
0
5
10
15
20
25
30
t/min

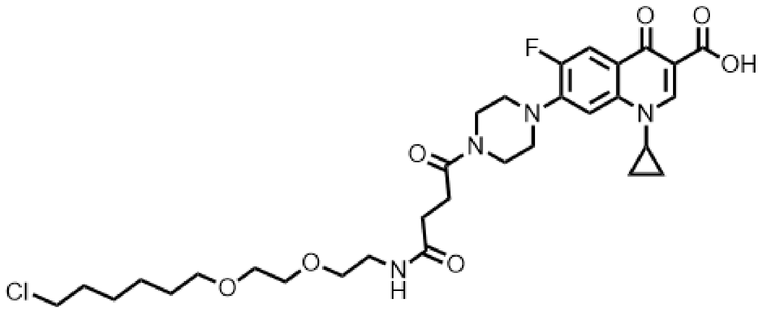
F
O
O
OH
N
N
N
O
O
N
H
O
O
Cl

(**6**)

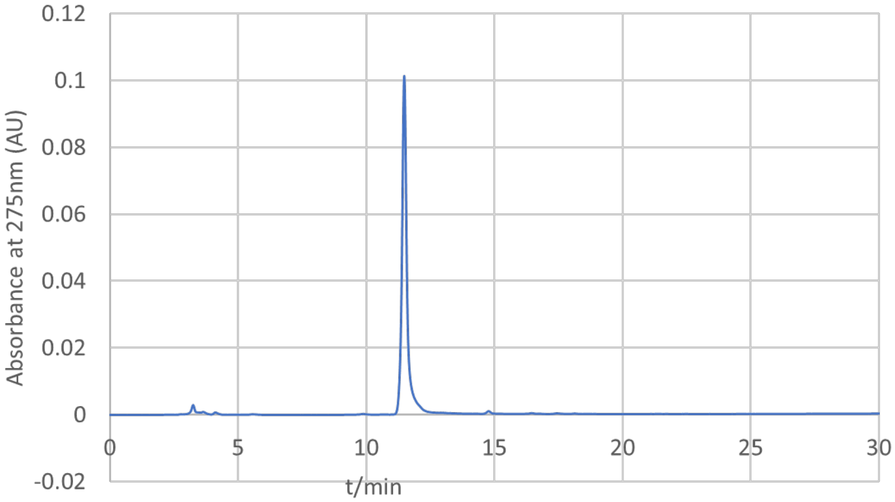
0.12
0.1
0.08
0.06
0.04
0.02
0
-0.02
Absorbance at 275nm (AU)
0
5
10
15
20
25
30
t/min

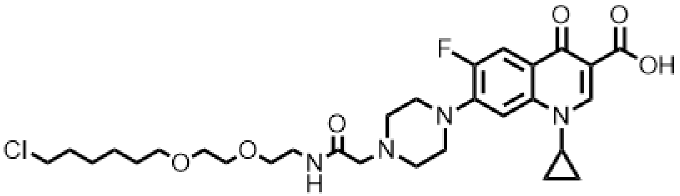
F
O
O
OH
N
N
N
Cl
O
O
N
H
O

(**7**)

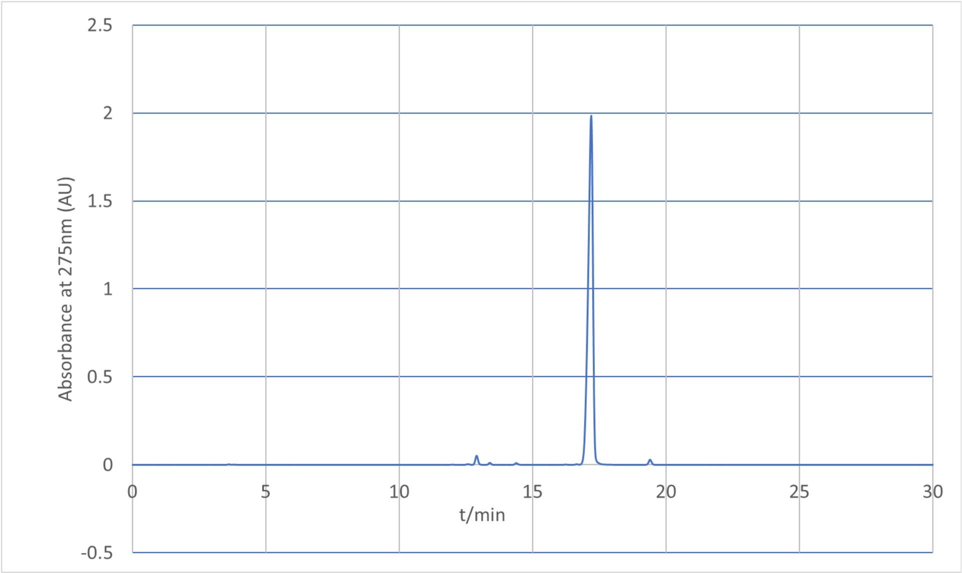
2.5
2
1.5
1
0.5
0
-0.5
Absorbance at 275nm (AU)
0
5
10
15
20
25
30
t/min

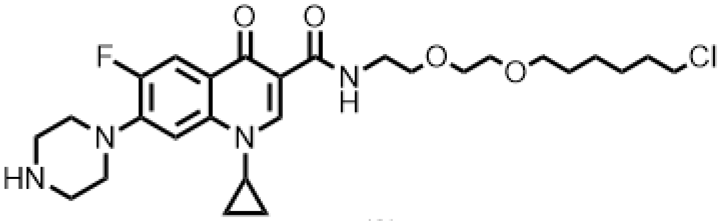
F
O
O
N
H
O
O
Cl
N
N
HN

(8)

Mass Spectra of Compounds:

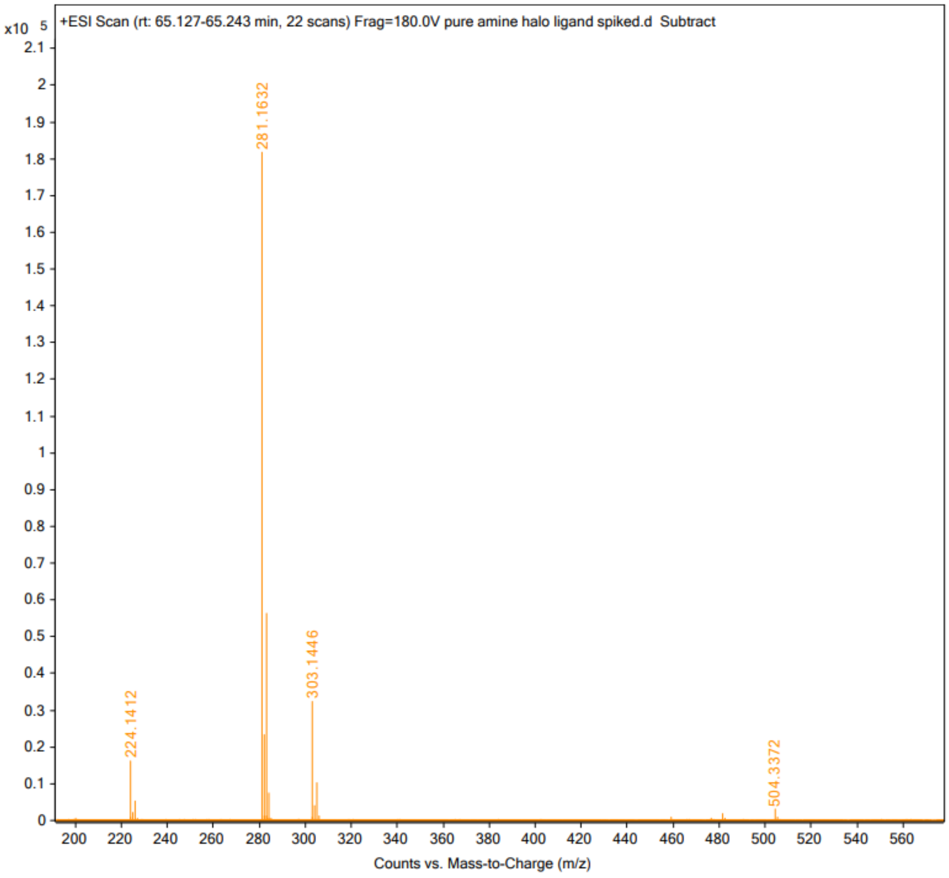


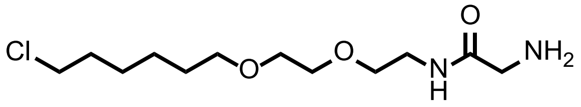


(**2**)

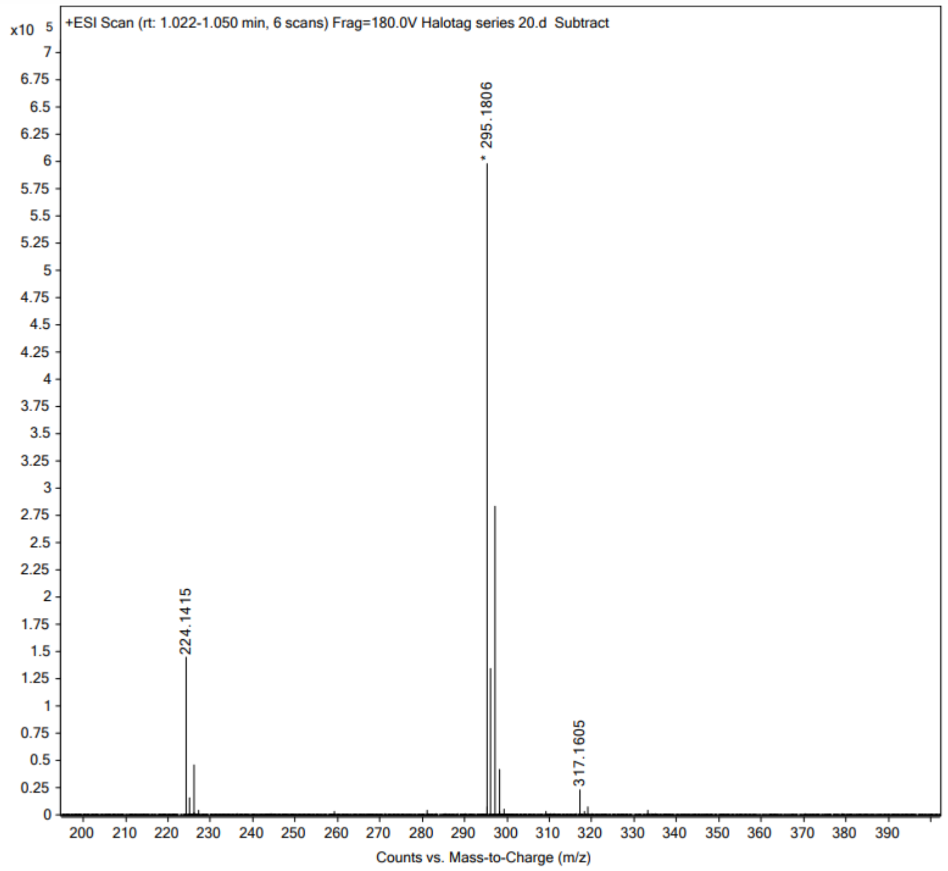

x10 5
+ESI Scan (rt: 1.022-1.050 min, 6 scans) Frag=180.0V Halotag series 20.d Subtract
* 295.1806
224.1415
317.1605
Counts vs. Mass-to-Charge (m/z)


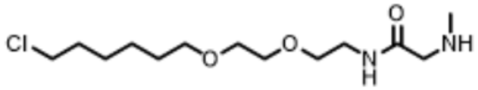

Cl
O
O
O
N
H
NH


(**3**)

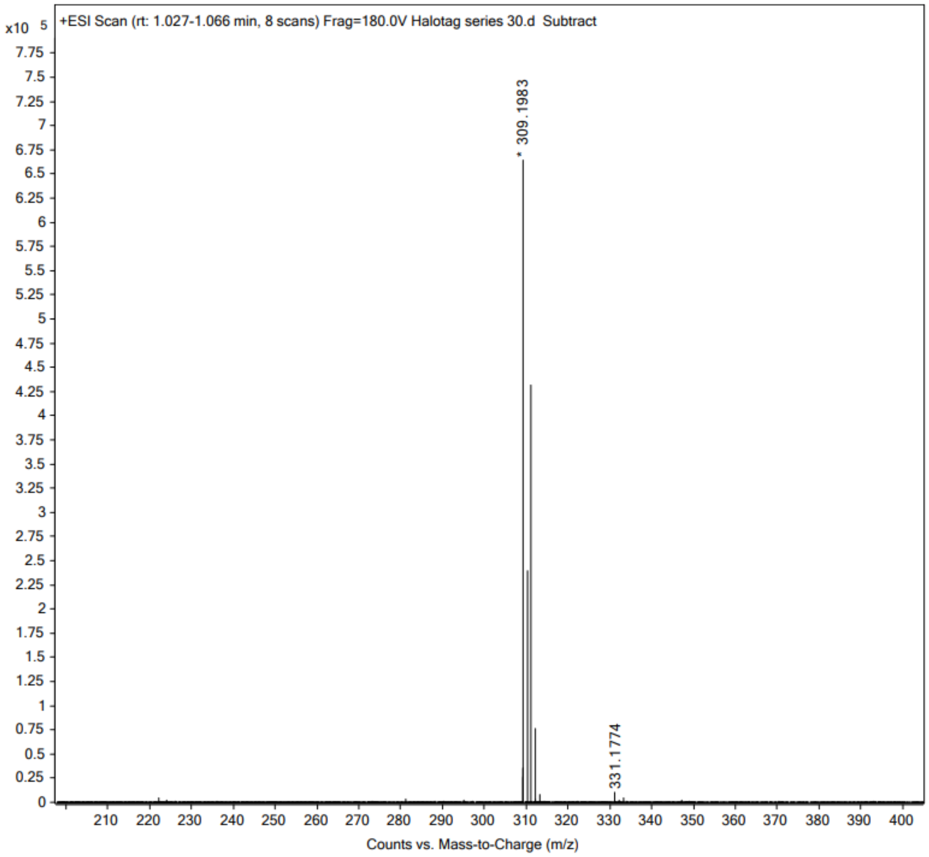

x10 5
+ESI Scan (rt: 1.027-1.066 min, 8 scans) Frag=180.0V Halotag series 30.d Subtract
* 309.1983
331.1774
Counts vs. Mass-to-Charge (m/z)


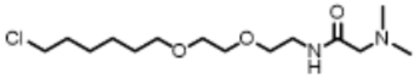

(4)

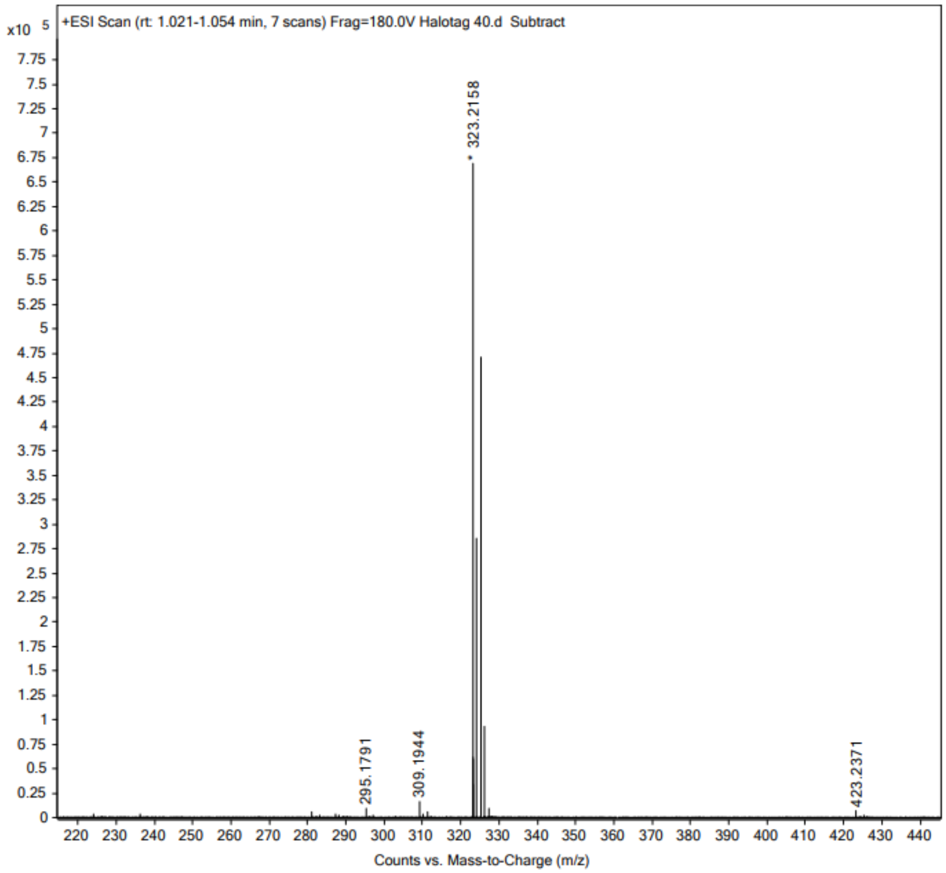
+ESI Scan (rt: 1.021-1.054 min, 7 scans) Frag=180.0V Halotag 40.d Subtract
x10 5
323.2158
295.1791
309.1944
423.2371
Counts vs. Mass-to-Charge (m/z)

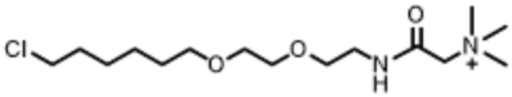
Cl
O
O
N
H
O
N
+

(**5**)

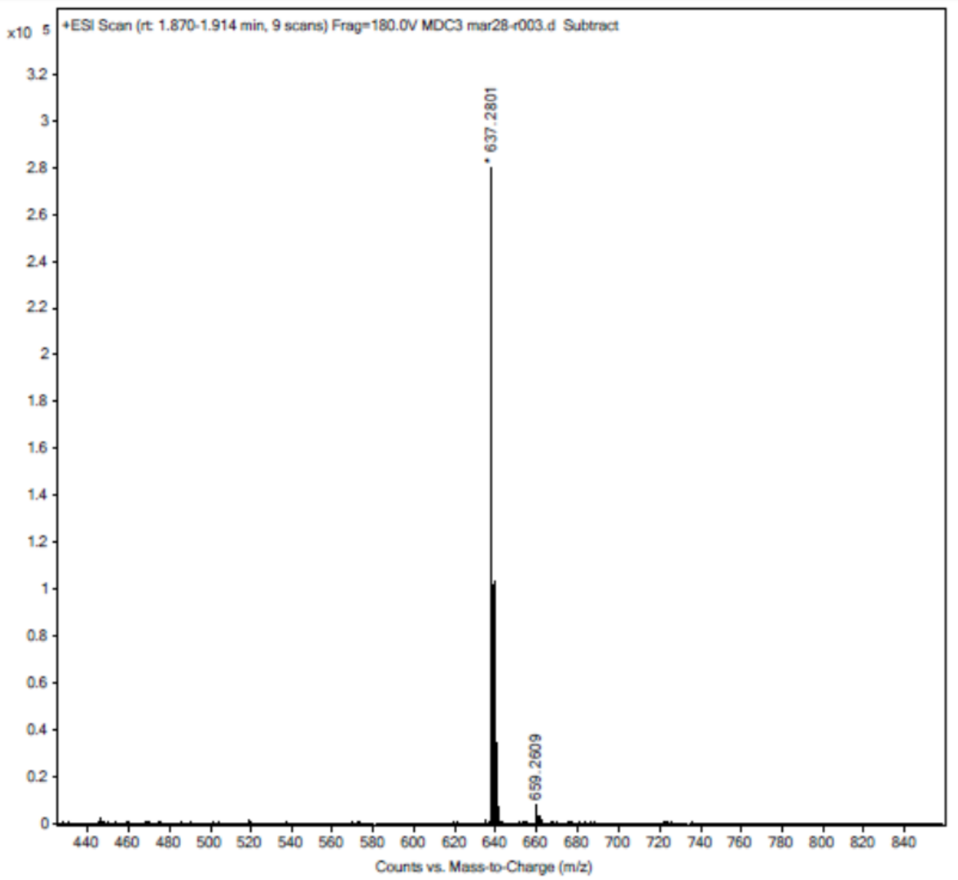

x10 5
+ESI Scan (rt: 1.870-1.914 min, 9 scans) Frag=180.0V MDC3 mar28-r003.d Subtract
* 637.2801
659.2609
Counts vs. Mass-to-Charge (m/z)


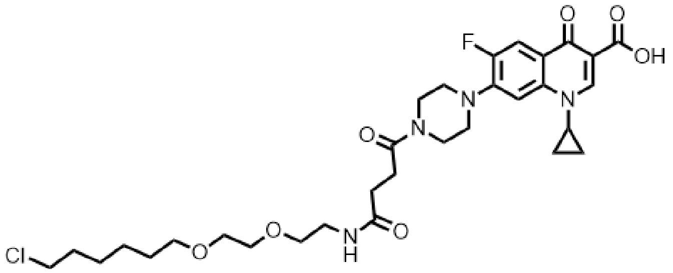

F
O
O
OH
N
N
N
O
O
N
H
O
O
Cl


(**6**)

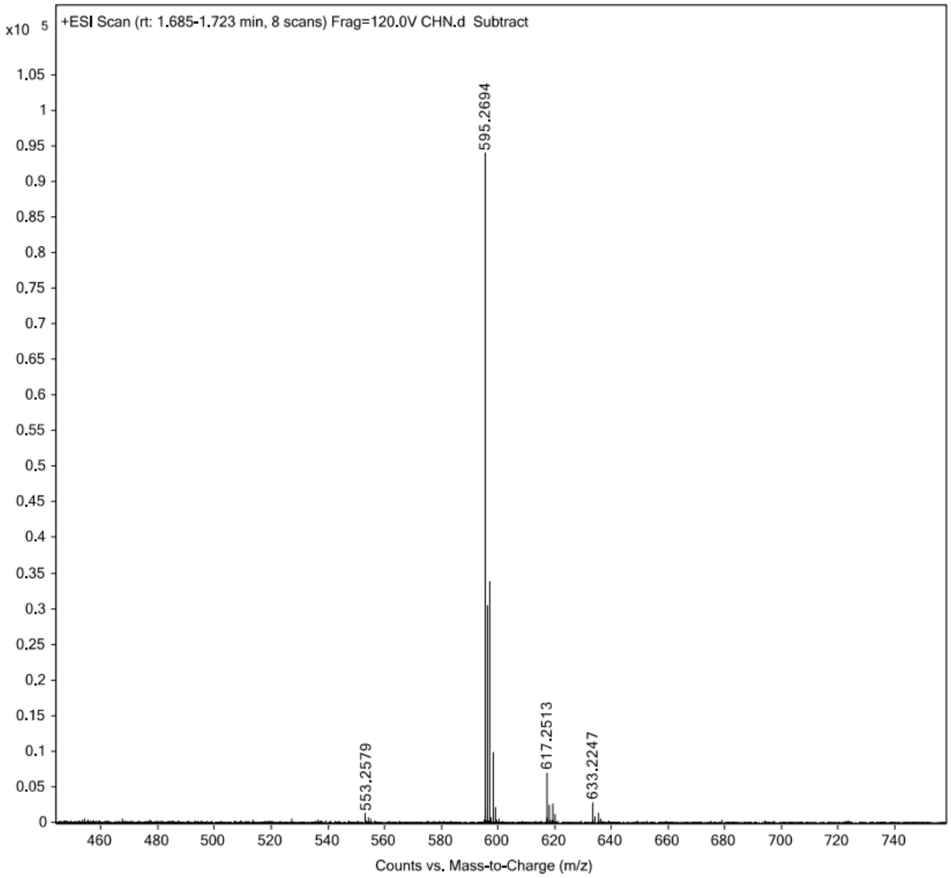

x10 5
+ESI Scan (rt: 1.685-1.723 min, 8 scans) Frag=120.0V CHN.d Subtract
595.2694
553.2579
617.2513
633.2247
Counts vs. Mass-to-Charge (m/z)


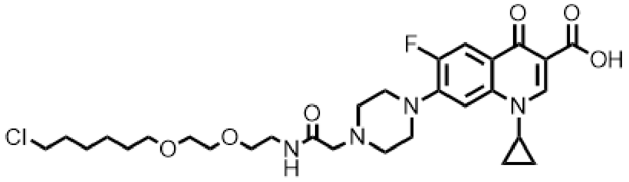

(**7**)

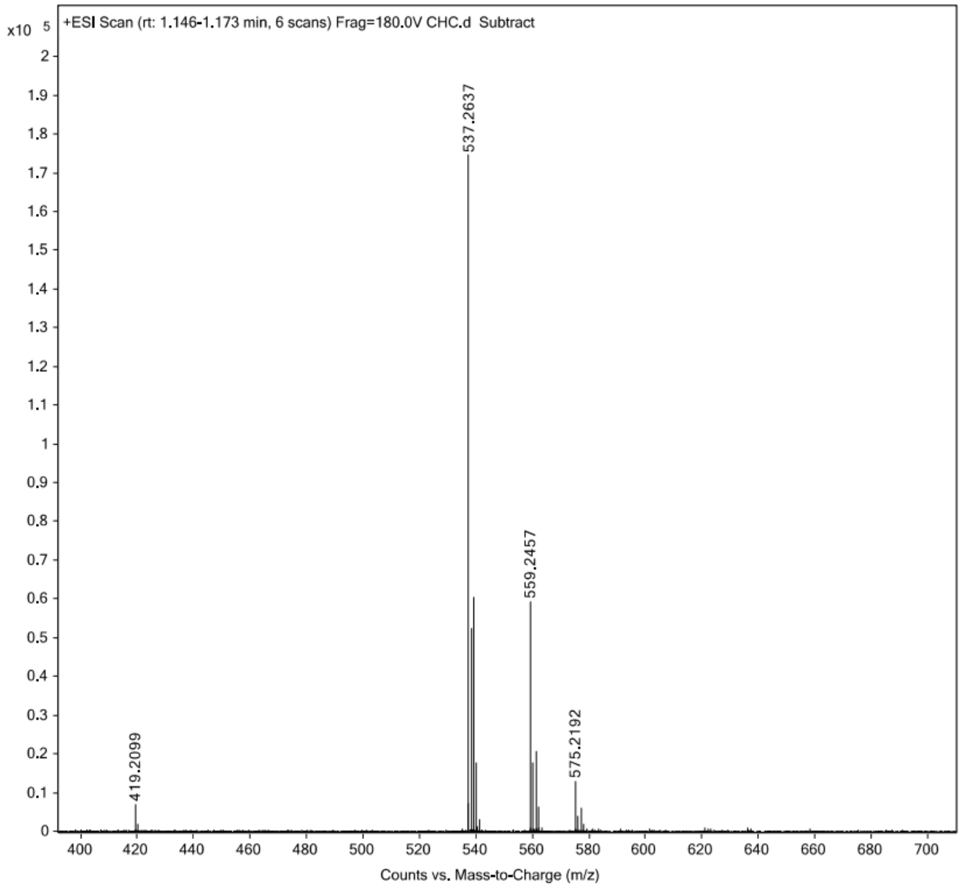

x10 5
+ESI Scan (rt: 1.146-1.173 min, 6 scans) Frag=180.0V CHC.d Subtract
537.2637
559.2457
575.2192
419.2099
Counts vs. Mass-to-Charge (m/z)


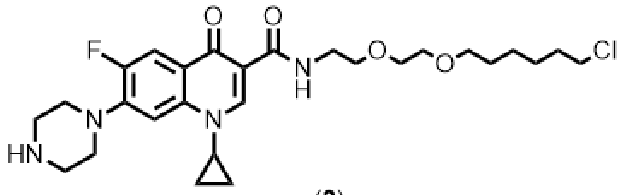

F
O
O
N
H
O
O
Cl
N
HN
N


(**8**)
